# Increased excitation-inhibition balance due to a loss of GABAergic synapses in the serine racemase knockout model of NMDA receptor hypofunction

**DOI:** 10.1101/2020.09.18.304170

**Authors:** Shekib A. Jami, Scott Cameron, Jonathan M. Wong, Emily R. Daly, A. Kimberley McAllister, John A. Gray

## Abstract

There is substantial evidence that both NMDA receptor (NMDAR) hypofunction and dysfunction of GABAergic neurotransmission contribute to schizophrenia, though the relationship between these pathophysiological processes remains largely unknown. While models using cell-type-specific genetic deletion of NMDARs have been informative, they display overly pronounced phenotypes extending beyond those of schizophrenia. Here, we used the serine racemase knockout (SRKO) mice, a model of reduced NMDAR activity rather than complete receptor elimination, to examine the link between NMDAR hypofunction and decreased GABAergic inhibition. The SRKO mice, in which there is a >90% reduction in the NMDAR co-agonist D-serine, exhibit many of the neurochemical and behavioral abnormalities observed in schizophrenia. We found a significant reduction in inhibitory synapses onto CA1 pyramidal neurons in the SRKO mice. This reduction increases the excitation/inhibition balance resulting in enhanced synaptically-driven neuronal excitability and elevated broad-spectrum oscillatory activity in *ex vivo* hippocampal slices. Consistently, significant reductions in inhibitory synapse density in CA1 were observed by immunohistochemistry. We further show, using a single-neuron genetic deletion approach, that the loss of GABAergic synapses onto pyramidal neurons observed in the SRKO mice is driven in a cell-autonomous manner following the deletion of SR in individual CA1 pyramidal cells. These results support a model whereby NMDAR hypofunction in pyramidal cells disrupts GABAergic synapse development leading to disrupted feedback inhibition and impaired neuronal synchrony.

## Introduction

Schizophrenia is a devastating psychiatric disease characterized by psychosis along with profound cognitive and social impairments. One prominent and enduring model implicates hypofunction of *N*-methyl-D-aspartate receptors (NMDARs) in the broad symptomatology of schizophrenia (Javitt and Zukin, 1991; Jentsch et al., 1997; Kirihara et al., 2012; Nakazawa and Sapkota, 2020). For example, open channel NMDAR inhibitors, such as phencyclidine (PCP) and ketamine, induce schizophrenia-like symptoms in healthy subjects (Krystal et al., 1994; Lahti et al., 2001), and exacerbate both positive and negative symptoms in patients with schizophrenia (Lahti et al., 1995b; Malhotra et al., 1997; Lahti et al., 2001), supporting a shared mechanism between NMDAR dysfunction and schizophrenia pathophysiology. In addition, mice with low levels of the obligatory GluN1 subunit of NMDA receptor, so-called GluN1 hypomorphs, display behaviors and endophenotypes consistent with schizophrenia (Mohn et al., 1999; Duncan et al., 2002; Duncan et al., 2004; Fradley et al., 2005; Duncan et al., 2006; Moy et al., 2006; Bickel et al., 2008; Dzirasa et al., 2009; Halene et al., 2009; Ramsey, 2009; Saunders et al., 2012).

Another well-supported hypothesis states that schizophrenia arises from changes in the ratio of excitatory to inhibitory activity in the brain (E/I imbalance), specifically through downregulation of GABAergic inhibition, and may represent a point of overlap between schizophrenia and autism (Lewis et al., 2005; Sohal and Rubenstein, 2019). Decreases in GABAergic markers in schizophrenia have been consistently observed in postmortem tissue (Lewis et al., 1999; Lewis et al., 2004; Lewis et al., 2008; Gonzalez-Burgos et al., 2011; Stan and Lewis, 2012; Glausier and Lewis, 2017). Furthermore, decreased GABAergic signaling disrupts oscillatory activity in the brain – particularly gamma oscillations (Lodge et al., 2009) – that may be important for a variety of cognitive processes (Sohal, 2016) including perceptual binding (Singer and Gray, 1995), cognitive flexibility (Cho et al., 2015), and attention (Tiesinga et al., 2004; Kim et al., 2016).

In the present study, we evaluated E/I balance in a mouse model of NMDAR hypofunction associated with the knockout of serine racemase (SR), the biosynthetic enzyme for the NMDAR co-agonist D-serine (Wolosker et al., 1999; Coyle and Balu, 2018). In contrast to mouse models using broad genetic deletion of NMDARs which have phenotypes extending beyond the bounds of schizophrenia phenomenology (Nakazawa et al., 2017), similar to the NMDAR hypomorph mice which have a severe reduction in NMDAR expression (Barkus et al., 2012; Gandal et al., 2012; Moy et al., 2012), the SRKO mice provide a more subtle and potentially physiologically relevant model of NMDAR hypofunction (Coyle and Balu, 2018). Indeed, deficiency of D-serine and the subsequent hypofunction of NMDARs has been implicated in the pathophysiology of schizophrenia (Coyle, 2012). Genetic studies have suggested that SR, as well as the degradation enzyme D-amino acid oxidase (DAAO) and G72, an activator of DAAO, are putative risk genes for schizophrenia (Chumakov et al., 2002; Detera-Wadleigh and McMahon, 2006; Goltsov et al., 2006; Morita et al., 2007; Shi et al., 2008). In addition, D-serine levels in the CSF and serum are decreased in individuals with schizophrenia (Hashimoto et al., 2003; Bendikov et al., 2007) and supplementation of antipsychotics with D-serine improves negative and cognitive symptoms in patients with schizophrenia (Tsai et al., 1998; Heresco-Levy et al., 2005; Lane et al., 2005). Consistent with well-characterized hallmarks of schizophrenia, the SRKO mice have reductions in cortical and hippocampal dendritic complexity and spine density, reduced hippocampal volume (Balu et al., 2012; Balu et al., 2013), and impaired performance on cognitive tasks that can be improved with exogenous D-serine administration (Basu et al., 2009; DeVito et al., 2011; Balu et al., 2012; Balu and Coyle, 2014).

Here we show that SRKO mice also have a significant reduction in GABAergic synapses onto the soma and apical dendrites of CA1 pyramidal neurons. This reduction in inhibition increases the E/I ratio resulting in enhanced synaptically-driven neuronal excitability and elevated broad-band spectral power *in vitro.* Single neuron deletion of SR revealed that the loss of inhibitory synapses is driven cell-autonomously by the loss of SR in the pyramidal neurons, consistent with recent evidence that NMDARs on pyramidal neurons regulate GABAergic synapse development (Lu et al., 2013; Gu et al., 2016; Gu and Lu, 2018). These results support a model of pyramidal cell NMDAR hypofunction directly leading to GABAergic dysfunction.

## Materials and Methods

### Animals

The SRKO mice are derived from the floxed SR mice (SR^fl^), in which the first coding exon (exon 3) is flanked by loxP sites as described (Basu et al., 2009; Benneyworth et al., 2012) and are maintained on a C57Bl/6J background. Mice were group-housed in polycarbonate cages and maintained on a 12-hour light/dark cycle. Animals were given access to food and water ad libitum. The University of California Davis Institutional Animal Care and Use Committee approved all animal procedures.

### Slice Preparation

Male SR^fl^ (labeled as WT) and SRKO mice (2–3 months old) were deeply anesthetized with isoflurane, followed by cervical dislocation and decapitation. The brain was rapidly removed and submerged in ice-cold, oxygenated (95% O_2_/5% CO_2_) ACSF containing (in mm) as follows: 124 NaCl, 4 KCl, 25 NaHCO_3_, 1 NaH_2_PO_4_, 2 CaCl_2_, 1.2 MgSO_4_, and 10 glucose (Sigma-Aldrich). On a cold plate, the brain hemispheres were separated, blocked, and the hippocampi removed. For extracellular recordings, 350-400-μm-thick slices were cut using a McIlwain tissue chopper (Brinkman, Westbury, NY). For whole-cell recordings, a modified transverse 300 μm slices of dorsal hippocampus were prepared by performing a ~10° angle blocking cut of the dorsal portion of each cerebral hemisphere (Bischofberger et al., 2006) then mounting the cut side down on a Leica VT1200 vibratome in ice-cold, oxygenated (95% O_2_/5% CO_2_) ACSF. Slices were incubated (at 32°C) for 20 minutes and then maintained in submerged-type chambers that were continuously perfused (2–3 ml/min) with oxygenated (95% O_2_/5% CO_2_) ACSF at room temperature and allowed to recover for at least 1.5-2 h before recordings. Just prior to the start of experiments, slices were transferred to a submersion chamber on an upright Olympus microscope, perfused with warmed to 30.4°C using a temperature controller (Medical System Corp.) normal ACSF saturated with 95% O_2_/5% CO_2_. For oscillation studies, after 1.5-2 h recovery, slices were transferred into a model RC-27L submersion recording chamber (Warner Instruments LLC, Hamden, CT) that allows the tissue slice to be perfused from both above and below, and perfused at an increased flow rate of perfusion (10 ml/min) (Hajos et al., 2009) with 34°C normal ACSF saturated with 95% O_2_/5% CO_2_.

### Extracellular recordings

A bipolar, nichrome wire stimulating electrode (MicroProbes) was placed in *stratum radiatum* of the CA1 region and used to activate Schaffer collateral (SC)-CA1 synapses. For extracellular recordings, evoked fEPSPs (basal stimulation rate = 0.033 Hz) were recorded in *stratum radiatum* using borosilicate pipettes (Sutter Instruments, Novato, CA) filled with ACSF (resistance ranged from 5–10 MΩ). To determine response parameters of excitatory synapses, basal synaptic strength was determined by comparing the amplitudes of presynaptic fiber volleys and postsynaptic fEPSP slopes for responses elicited by different intensities of SC fiber stimulation. Presynaptic neurotransmitter release probability was compared by paired-pulse ratio (PPR) experiments, performed at 25, 50, 100, and 200 msec stimulation intervals. LTP was induced by high-frequency stimulation (HFS) using a 1x tetanus (1-s-long train of 100 Hz stimulation). At the start of each experiment, the maximal fEPSP amplitude was determined and the intensity of presynaptic fiber stimulation was adjusted to evoke fEPSPs with an amplitude ~30-40% of the maximal amplitude. The average slope of EPSPs elicited 55–60 min after HFS (normalized to baseline) was used for statistical comparisons. For experiments performed in picrotoxin (PTX, Sigma-Aldrich; 50 μM) the CA3 region was removed. For oscillation induction experiments, local field potentials were recorded with a borosilicate pipette (Sutter Instruments, Novato, CA) filled with ACSF (resistance ranged from 8-10 MΩ) placed into the CA1 pyramidal cell layer. Signals were low-pass filtered at 3 kHz, high-pass filtered at 0.3 Hz, and digitized at 20 kHz for off-line analysis. Gamma oscillations were induced by bath-application of carbachol (CCh, 25 μM) (Fellous and Sejnowski, 2000) after 3-5 min of baseline recording. For power spectrum analyses the data were filtered at 5 Hz-200 Hz with a band-pass eight-pole Bessel filter. Mainline noise (60 Hz and 120 Hz) was eliminated manually. Analyses were performed with the Clampex 10.6 software suite (Molecular Devices, San Jose, CA) and Prism 8.0 software (GraphPad Software, San Diego, CA).

### Whole-cell current clamp recordings

CA1 pyramidal neurons were visualized by infrared differential interference contrast microscopy, and current clamp recordings were performed using borosilicate recording electrodes (3–5 MΩ) filled with a K^+^-based electrode-filling solution containing (in mM)-135 K-gluconate, 5 NaCl, 10 HEPES, 2 MgCl, 0.2 EGTA, 10 Na_2_-phosphocreatine, 4 Na-ATP, 0.4 Na-GTP (pH = 7.3, 290 mOsm). Passive and active membrane properties of CA1 pyramidal cells were determined using three 500 ms current pulses 10 s apart. Current injections were first recorded in increasing order (i.e. 0, 25, 50, 75, 100, 125, 150, and 200 pA) and then in decreasing order. Values obtained from the responses elicited by the same current injection were averaged. For input resistance, 500 ms current steps of 0 to −200 pA were injected in −20 pA increments. Steady-state responses were measured as the average change in voltage in the last 100 ms of the pulse. The slope of a regression line fitted to the voltage versus current data was used to calculate input resistance. Sag currents were measured during the 100 pA hyperpolarizing steps and calculated as the initial voltage trough minus the steady-state voltage change. Firing frequency versus injected current was measured as the number of spikes per 500 ms step in 25 pA increments from 0 to 200 pA. Rheobase was determined by injecting 0.5 ms square pulses in 2 pA steps and recording the strength of the first pulse to elicit an action potential. Spike firing threshold and AP height were calculated by injecting a 2 ms square pulse of 1.8 nA. The medium afterhyperpolarization (mAHP) was somatically elicited with a 50 Hz train of suprathreshold (1.8 nA) current injections. After zeroing the baseline, the mAHP was measured as the negative peak after the membrane potential recrossed zero after the last action potential. To measure the E/I ratio from CA1 pyramidal neurons, current clamp recordings at resting membrane potential (RMP) were made in the absence of synaptic blockers. Stimulus strength was adjusted to evoke both monosynaptic EPSPs and a compound di-synaptic/ monosynaptic IPSPs that was 40-60% of the maximum inhibitory component (Bateup et al., 2013; Bartley et al., 2015). E/I ratio was calculated from averaged baseline subtracted traces as the maximum depolarization amplitude (in mV) divided by the maximum hyperpolarization amplitude in the 300 ms after the stimulus. Synaptically-mediated excitability was determined with short trains of synaptic stimulation (5 pulses at 100 Hz SC fiber stimulation) with the CA1 pyramidal neurons at RMP in the absence of synaptic blockers. Stimulation strength was adjusted so that the initial EPSP was ~5 mV for each recording.

### Whole-cell voltage clamp recordings

CA1 pyramidal neurons were visualized by infrared differential interference contrast microscopy, and voltage-clamp recordings were performed using borosilicate glass recording pipettes (3–5 MΩ) filled with a Cs^+^-based electrode-filling solution containing (in mM): 135 Cs-methanesulfonate, 8 NaCl, 5 QX314 (Sigma-Aldrich), 0.3 EGTA, 4 Mg-ATP, 0.3 Na-GTP, and 10 HEPES (pH = 7.3, 290 mOsm). Evoked IPSCs (eIPSCs), spontaneous IPSCs (sIPSCs), and miniature IPSCs (mIPSCs) were recorded in presence of APV and NBQX (Tocris; 50 μM and 10 μM APV respectively) to block AMPAR and NMDAR currents. Miniature EPSCs (mEPSCs) and mIPSCs were also recorded in the presence of 0.5 μM tetrodotoxin (TTX; Alomone Laboratories, Jerusalem, Israel) to block action potential-dependent neurotransmitter release. The outward IPSCs were completely blocked by PTX (50 μM). For the input/output (I/O) curves of eIPSCs, the stimulus intensity of the threshold evoked response was first determined and then stimulation was increased to develop the I/O curves. Recordings where series resistance was ≥ 25 MΩ or unstable were discarded. Series resistance compensation was used in all voltage-clamp recordings except in experiments examining miniature postsynaptic currents. All recordings were obtained with a MultiClamp 700B amplifier (Molecular Devices), filtered at 2 kHz, digitized at 10 Hz. Analysis was performed with the Clampex 10.6 software suite and GraphPad Prism 8.0.

### Single neuron SR deletion experiments

Neonatal [P0-P1] SR^fl^ mice of both sexes were stereotaxically injected with high-titer rAAV1-Cre:GFP viral stock (~1-5 x 10^12^ vg/mL) with coordinates targeting hippocampal CA1 as previously described (Gray et al., 2011; Wong and Gray, 2018). At 2-3 months, the injected mice were anesthetized with isoflurane and transcardially perfused with ice-cold artificial cerebrospinal fluid (ACSF), containing (in mM) 119 NaCl, 26.2 NaHCO_3_, 11 glucose, 2.5 KCl, 1 NaH_2_PO_4_, 2.5 CaCl_2_, and 1.3 MgSO_4_. Modified transverse 300 μm slices of dorsal hippocampus were prepared by performing a ~10° angle blocking cut of the dorsal portion of each cerebral hemisphere (Bischofberger et al., 2006) then mounting the cut side down on a Leica VT1200 vibratome in ice-cold cutting buffer. Slices were incubated in 32°C NMDG solution containing (in mM) 93 NMDG, 93 HCl, 2.5 KCl, 1.2 NaH_2_PO_4_, 30 NaHCO_3_, 20 HEPES, 25 glucose, 5 sodium ascorbate, 2 thiourea, 3 sodium pyruvate, 10 MgSO_4_, and 0.5 CaCl_2_ (Ting et al., 2018) for 15 mins, transferred to room temperature ACSF and held for at least 1 hr before recording. All solutions were vigorously perfused with 95% O_2_ and 5% CO_2_. Slices were transferred to a submersion chamber on an upright Olympus microscope, perfused in room temperature ACSF, and saturated with 95% O_2_ and 5% CO_2_. CA1 neurons were visualized by infrared differential interference contrast microscopy, and GFP+ cells were identified by epifluorescence microscopy. Cre expression was generally limited to the hippocampus within a sparse population of CA1 pyramidal neurons. Cells were patched with 3–5 MΩ borosilicate pipettes filled with intracellular solution containing (in mM) 135 cesium methanesulfonate, 8 NaCl, 10 HEPES, 0.3 Na-GTP, 4 Mg-ATP, 0.3 EGTA, and 5 QX-314 (Sigma, St Louis, MO) and mIPSCs were recorded at 0 mV in the presence of 50 μM APV, 10 μM NBQX, and 0.5 μM TTX. Series resistance was monitored and not compensated, and cells were discarded if series resistance varied more than 25%. Recordings were obtained with a Multiclamp 700B amplifier (Molecular Devices, Sunnyvale, CA), filtered at 2 kHz, and digitized at 10 Hz. Analysis was performed with the Clampex 10.6, MiniAnalysis, and GraphPad Prism 8.0 (GraphPad Software, San Diego, CA, USA).

### Immunohistochemistry

Male C57Bl/6J, SR^fl^ (labeled as WT) and SRKO mice (2–3 months old) were deeply anesthetized with isoflurane and injected with a lethal dose of Fatal Plus (Vortech Pharmaceuticals) pentobarbital solution. The mice were then perfused transcardially with 1xPBS followed by 4% paraformaldehyde (PFA; Electron Microscopy Sciences) in 1xPBS. Brains were removed and post-fixed for 3 h in 4% PFA in 1xPBS. The fixed brains were then cryoprotected stepwise, first in 10% sucrose in 1xPBS overnight, then in 30% sucrose in 1xPBS overnight. Brains were then mounted and frozen in O.C.T. compound (Tissue-Tek®). Coronal sections through the dorsal hippocampus were cut on a Leica CM3050 S cryostat at 10 μm and collected onto Superfrost™ Plus slides (Fisher). Sections were outlined with a hydrophobic barrier pen and all subsequent incubation steps were performed in a humidified chamber. The sections were blocked with 10% normal donkey serum in 1xPBS-T (0.5% Triton X-100) for 1 h at room temperature and then probed overnight with rabbit anti-VGAT antibody (Synaptic Systems, cat# 131 003, 1:500) in blocking solution at 4°C. The next day sections were rinsed 3x with 1xPBS-T and then incubated with secondary antibody (Donkey anti-rabbit 647, Jackson, cat# 711-605-152, 1:400) in 1xPBS-T for 90 mins at room temperature. The sections were then rinsed 3x with 1xPBS-T and counterstained with DAPI. The sections were then mounted with Mowoil® mounting medium and covered with a glass coverslip. After drying a series of images covering the hippocampus were collected on a Nikon C2 LSM with a Nikon CFI Apo Lambda 60x 1.4 NA oil objective. Laser and PMT settings remained constant between individuals and genotypes. Single images covering the regions of interest were stitched in together in Nikon Elements software. Regions of interest of the *stratum pyramidale* or *stratum radiatum* of hippocampal CA1 were analyzed using custom-written journals (Elmer et al., 2013) in Metamorph software (v7.5, Molecular Devices) to identify and quantify VGAT puncta density and intensity. Two regions of interest for both *stratum pyramidale* or *stratum radiatum* were analyzed for each of 3 individual animals per genotype. Data were graphed and analyzed using GraphPad Prism 8.0 (GraphPad Software, San Diego, CA, USA). Unpaired student’s t-tests were used to test for statistically significant differences between genotypes.

### Statistical analysis

Statistical comparisons were made with Student’s unpaired t-test or two-way ANOVA with Bonferroni’s multiple comparisons test as specified and appropriate, using GraphPad Prism 8.0 (GraphPad Software, San Diego, CA, USA). Spontaneous and miniature inhibitory synaptic events were analyzed using Mini Analysis software (Synaptosoft, Fort Lee, NJ, USA). Peaks of events were first automatically detected by the software according to a set threshold amplitude of 6 pA. To generate cumulative probability plots for both amplitude and inter-event time interval, the same number of events (50–200 events acquired after an initial 3 min of recording) from each CA1 pyramidal neuron was pooled for each group, and input into the Mini Analysis program. The Kolmogorov-Smirnov two-sample statistical test (KS test) was used to compare the distribution of events between WT and SRKO mice. For power spectrum analyses the data were filtered at 5 Hz-200 Hz with a band-pass eight-pole Bessel filter.

## Results

### Increased E/I balance in CA1 pyramidal cells in SRKO mice

To investigate the properties of excitatory synaptic transmission in the SRKO mice, we first conducted extracellular field recordings of SC-CA1 synapses in the SRKO mice. Consistent with previous studies (Balu et al., 2013), the basal excitatory synaptic strength, determined by comparing the amplitudes of presynaptic fiber volleys and field EPSP (fEPSP) slopes for responses elicited by different intensities of SC fiber stimulation (input-output curve), was unaltered in SRKO compared to WT slices (**Fig. 1A**, p=0.49, two-way ANOVA, F(1,122)=0.467). There was also no change in paired-pulse ratio (PPR) of the fEPSPs in the SRKO mice compared with WT mice (**Fig. 1B**, p=0.91, two-way ANOVA, F(1,23)=0.012), together suggesting no alteration of excitatory neurotransmission or presynaptic glutamate release probability in the SRKO mice. We next examined the input-output (I/O) function of evoked IPSCs (eIPSCs) which indicated a significant decrease in synaptic inhibition in SRKO mice compared with WT mice (**Fig. 1C**, p=0.0001, two-way ANOVA, F(1,224)=58.90; Bonferroni’s multiple comparisons test, *p<0.05). This difference was characterized by a downward shift in the I/O curve showing the relationship between eIPSC amplitude and stimulus intensity. PPR of eIPSCs was also unchanged in SRKO mice compared to WT mice (**Fig. 1D**, p=0.82, unpaired t-test, t(34)=0.230), suggesting that the reduction in inhibitory currents was not due to a change in the probability of GABA release. Using whole-cell current clamp recordings, we next examined the impact of the reduction in synaptic inhibition in the SRKO on the E/I ratio in CA1 pyramidal neurons by recording compound EPSP/IPSPs at RMP upon SC stimulation. We found no significant change in the amplitude of the EPSP component of the compound EPSP/IPSP (**Fig. 1E**, peak EPSP amplitude, p=0.07, unpaired t-test, t(24)=0.347), but there was a significant reduction in the IPSP component (**Fig. 1E**, peak IPSP amplitude, p=0.045, unpaired t-test, t(24)=2.11). This decrease in IPSP amplitude results in an increased E/I ratio (**Fig. 1E**, E/I ratio, p=0.035, unpaired t-test, t(24)=2.23). Together, these results suggest that a surprisingly selective GABAergic impairment in the SRKO mice leads to an increase in the E/I balance.

**Figure 1:**
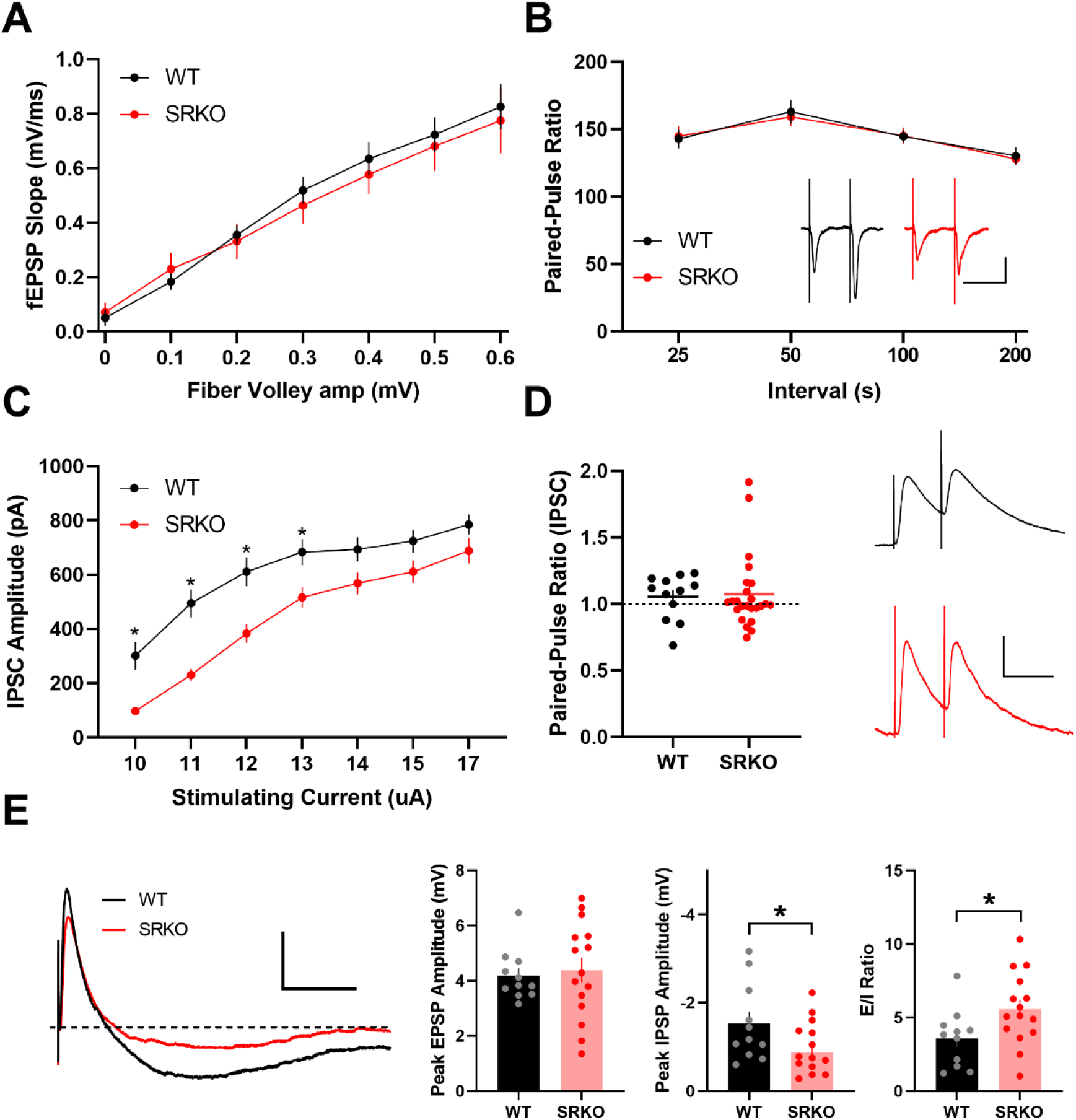
Increased E/I ratio in SRKO mice **(A)** Normal basal synaptic transmission as measured by presynaptic fiber volley amplitudes and postsynaptic fEPSP slopes for responses elicited by different intensities of SC fiber stimulation in WT (n=10) and SRKO (n=10) hippocampal slices (p=0.49, two-way ANOVA, F(1,122)=0.467). **(B)** Paired-pulse ratio is unchanged at SRKO SC-CA1 synapses compared to WT (p=0.91, two-way ANOVA, F(1,23)=0.012; WT: n=12, SRKO: n=13). Right, traces represent fEPSPs evoked by stimulation pulses delivered with a 50 ms interpulse interval; scale bars: 0.5 mV, 50 ms. **(C)** Input-output function of evoked IPSC amplitude versus stimulating current strength show a significant decrease in inhibition in SRKO mice (p=0.0001, two-way ANOVA, F(1,224)=58.90; Bonferroni’s multiple comparisons test, *p<0.05; WT n=17, SRKO n=17). Representative traces of evoked IPSC from WT and SRKO CA1 pyramidal cells at holding potential of 0 mV. **(D)** Paired pulse ratio of IPSCs plotted as a function of 50 ms ISI (WT: 1.056±0.049, n=12; SRKO: 1.076±0.057, n=24) indicating that there is no change in probability of inhibitory neurotransmitter release from presynaptic terminals. Representative traces of evoked IPSCs from WT and SRKO CA1 pyramidal cells; scale bars: 50 pA, 50 msec. Error bars show ± SEM. **(E)** Left, overlaid traces of compound excitatory (EPSP) and inhibitory (IPSP) postsynaptic potentials evoked by Schaffer collateral stimulation in absence of synaptic blockers at RMP from SRKO (red) and WT (black) mice; dashed line indicates the baseline; scale bars: 2 mV, 100 msec. Peak EPSP amplitude is unchanged between SRKO and WT mice. (WT: 4.2±0.3 mV, n=11; SRKO: 4.4±0.5, n=15). Peak IPSP amplitude is significantly decreased in SRKO mice compared to WT mice (WT: 1.5±0.3 mV, n=11; SRKO: 1.0±0.1, n=15). The E/I ratio in CA1 pyramidal cells calculated from EPSP and IPSP peak amplitudes is greater in SRKO mice compared to WT (WT: 3.6±0.6, n=11; SRKO: 5.6±0.6, n=15). Data represent mean ± SEM.

### Enhanced pyramidal cell excitability to synaptic stimulation in SRKO mice

Synaptic inhibition plays a key role in synaptic integration and spike initiation in neurons (Gulledge et al., 2005). Indeed, at hippocampal SC-CA1 synapses, EPSP-spike potentiation, an enhancement of spike probability in response to a synaptic input of a fixed slope, is dependent on changes in GABAergic inhibition (Marder and Buonomano, 2004). Thus, in the SRKO mice, we examined EPSP-spike coupling using short trains of SC stimulation (5 pulses at 100 Hz). Stimulation intensity was adjusted for each neuron to normalize the initial subthreshold EPSP to ~5 mV. We found a significantly increased probability of spiking in SRKO CA1 pyramidal cells compared to WT (**Fig. 2A**, p=0.0002, two-way ANOVA, F(1,115)=14.39), especially for the second stimulus (*p=0.013, Bonferroni’s multiple comparisons test, F(115)=3.07). Importantly, there were no differences in the intrinsic excitability of CA1 pyramidal cells between SRKO and WT mice **(Fig. 2B, Table 1)**. We analyzed the number of spikes elicited during 500 ms steps of somatically injected current and found no significant differences in the number of spikes between WT and SRKO neurons at steps of any intensity (**Fig. 2B**, p=0.759, two-way ANOVA, F(8,207)=0.621). There were also no significant differences in the passive membrane properties, rheobase, action potential threshold or height, or sag amplitude between the CA1 neurons of WT and SRKO mice **(Table 1)**. There was, however, a significant decrease in the amplitude of the post-burst AHP peak in the SRKO mice compared to WT (**Table 1**, p=0.028, unpaired t-test, t(23)=2.35). To investigate how the increase in E/I balance in the SRKO mice affects hippocampal network activity, we bath applied cholinergic agonist carbachol (25 μM) to *ex vivo* hippocampal slices to induce oscillatory activity in the slice (Williams and Kauer, 1997). We found extracellular field potential power was increased across the entire measured spectrum in slices from the SRKO mice compared with WT mice **(Fig. 2C)**, particularly in the slow gamma range of 30-50Hz (**Fig. 2C**, p<0.0001, two-way ANOVA, F(34,17782)=2067), where carbachol-induced gamma oscillations are commonly observed in ex vivo slices (Fisahn et al., 1998). Together, these data suggest that a reduction in inhibitory input onto CA1 pyramidal neurons in the SRKO mice increases the E/I balance resulting in enhanced synaptically-driven neuronal excitability leading to broad-band increases in carbachol-induced network activity in hippocampal slices.

**Figure 2:**
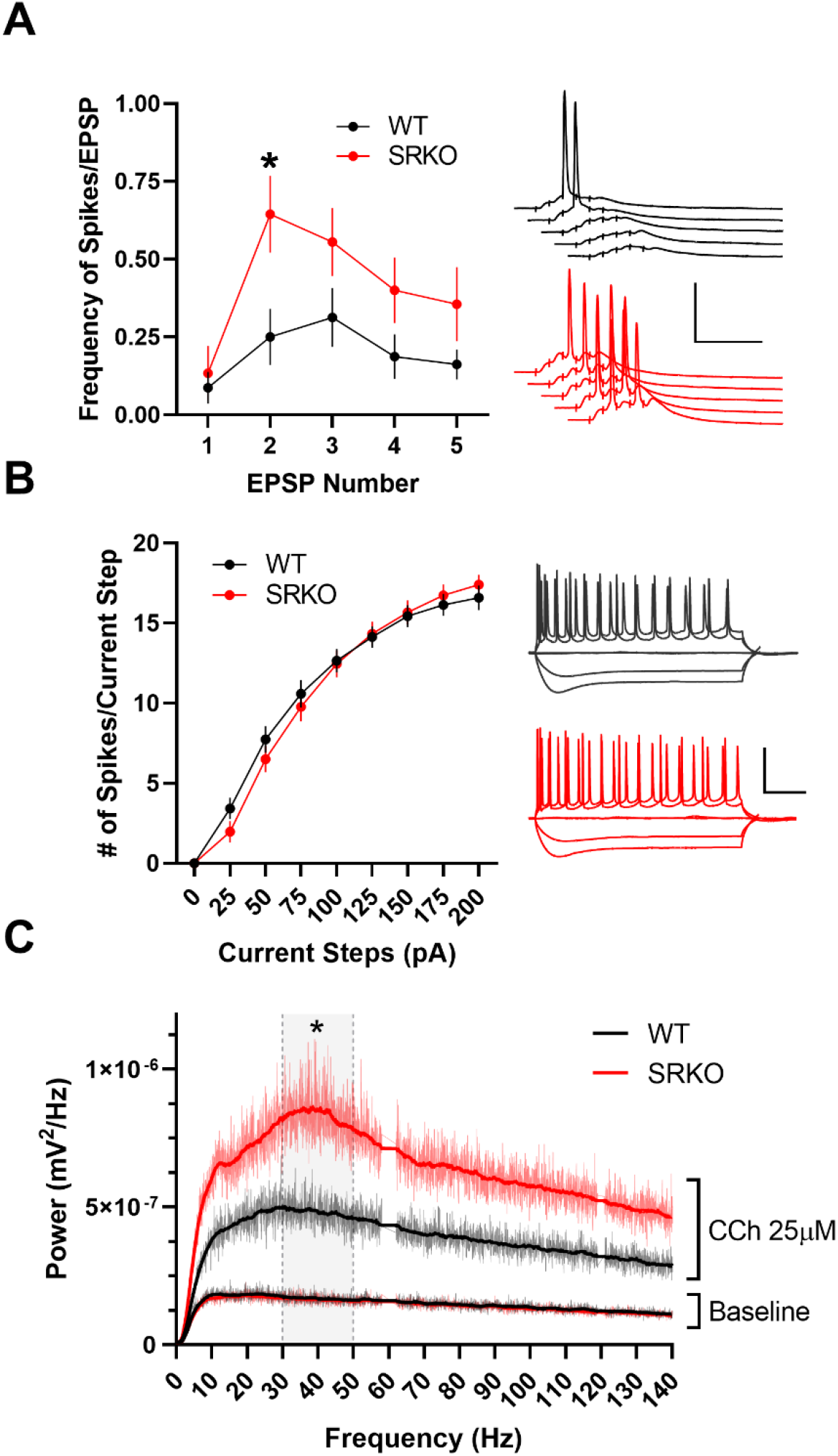
Increased synaptic excitability in SRKO mice **(A)** Short trains of synaptic stimulation leads to significantly more APs/EPSP in SRKO compared to WT (p=0.013, two-way ANOVA, F(115)=3.07; Bonferroni’s multiple comparisons test, *p<0.05); WT: n=16, SRKO: n=9). Right, sample traces of APs/EPSPs evoked by 5 pulses at 100 Hz SC fiber stimulation; scale bars: 50 mV, 50 msec. **(B)** Intrinsic excitability is unchanged in SRKO CA1 pyramidal neurons. Depolarization induced by somatic current injection elicits similar numbers of APs in WT and SRKO cells (p=0.759, two-way ANOVA, F(8,207)=0.6212; WT: n=12, SRKO: n=13) suggesting basal synaptic transmission is unaffected. Sample traces for 0, −100, −200, +100, and +200 pA current steps; scale bars: 50 mV, 100 msec. Data represent mean ± SEM. **(C)** Mean power spectrum of carbachol-induced hippocampal CA1 oscillations in SRKO and WT mice. There is a statistically significant increase in power between 30 and 50 Hz (p<0.0001, two-way ANOVA, F(34,17782)=2067), whereas the spectra obtained during the baseline period appear as nearly overlapping flat lines. Main line noise (60 Hz and 120 Hz) was eliminated manually. Solid lines are derived from a smoothing function.

**Table 1:** Intrinsic Excitability in Wild-Type and SRKO CA1 Pyramidal Neurons

| Property | WT<br>(n=12) | SRKO<br>(n=13) | Unpaired<br>t-test | P<br>value |
| --- | --- | --- | --- | --- |
| RMP (mV) | 59.2 ± 1.1<br>[-61 to -53] | -60.4 ± 0.7<br>[-64 to -55] | t(23)=0.01 | 0.207 |
| R<sub>input</sub>(MΩ) | 174.6 ± 13.0<br>[97.7 – 235.8] | 154.0 ± 9.9<br>[89.4 – 219.3] | t(23)=1.27 | 0.216 |
| Sag (mV)<sup>A</sup> | -0.20 ± 0.001<br>[-0.17 to -0.24] | -0.22 ± 0.01<br>[-0.14 to -0.26] | t(23)=1.27 | 0.219 |
| Rheobase (pA) | 15.1 ± 5.1<br>[2.7 – 59.3] | 25.2 ± 4.8<br>[2.7 – 70.2] | t(23)=1.45 | 0.159 |
| AP Threshold (mV)<sup>A</sup> | 49.2 ± 0.8<br>[-44.3 to -52.7] | -50.4 ± 0.6<br>[-44.8 to -53.1] | t(23)=1.19 | 0.247 |
| AP Height (mV) | 120.9 ± 2.3<br>[105 to 140] | 120.9 ± 1.7<br>[110 to 134] | t(23)=0.01 | 0.989 |
| AHP Peak (mV) | -2.79 ± 0.4<br>[-5.75 to -0.845] | -1.60 ± 0.3<br>[-3.74 to -0.46] | t(23)=2.35 | 0.028* |

Mean ± SEM [range]
<sup>A</sup> junction potential not adjusted

### Loss of picrotoxin-induced disinhibition during LTP in SRKO mice

In hippocampal SC-CA1 field LTP experiments induced with a HFS (e.g. 100 Hz tetanus), the addition of a GABA_A_ inhibitor (e.g. PTX) causes a disinhibition that enhances LTP (**Fig. 3A**, p=0.0002, unpaired t-test, t(26)=4.38) (Wigstrom and Gustafsson, 1983). Due to the reduced inhibition observed in the SRKO mice, we hypothesized that PTX-induced disinhibition might be disrupted. Consistently, we found that, in hippocampal slices from the SRKO mice, the addition of PTX (50 μM) did not affect the magnitude of LTP induced with a single 100 Hz tetanus (**Fig. 3B**, p=0.394, unpaired t-test, t(21)=0.871). Interestingly, comparing data between WT and SRKO slices, we only observed significantly different LTP in the presence of PTX (p=0.046, unpaired t-test, t(22)=2.11). In the absence of PTX, there was no difference in LTP between WT and SRKO slices (p=0.623, unpaired t-test, t(25)=0.498), likely due to baseline disinhibition in the SRKO slices. Thus, by removing the impact of the reduced inhibition in the SRKO slices, the addition of PTX provides a more direct measure of the impact of synaptic NMDAR hypofunction in LTP, consistent with previous studies (Basu et al., 2009; Henneberger et al., 2010; Benneyworth et al., 2012; Balu et al., 2013; Balu et al., 2016).

**Figure 3:**
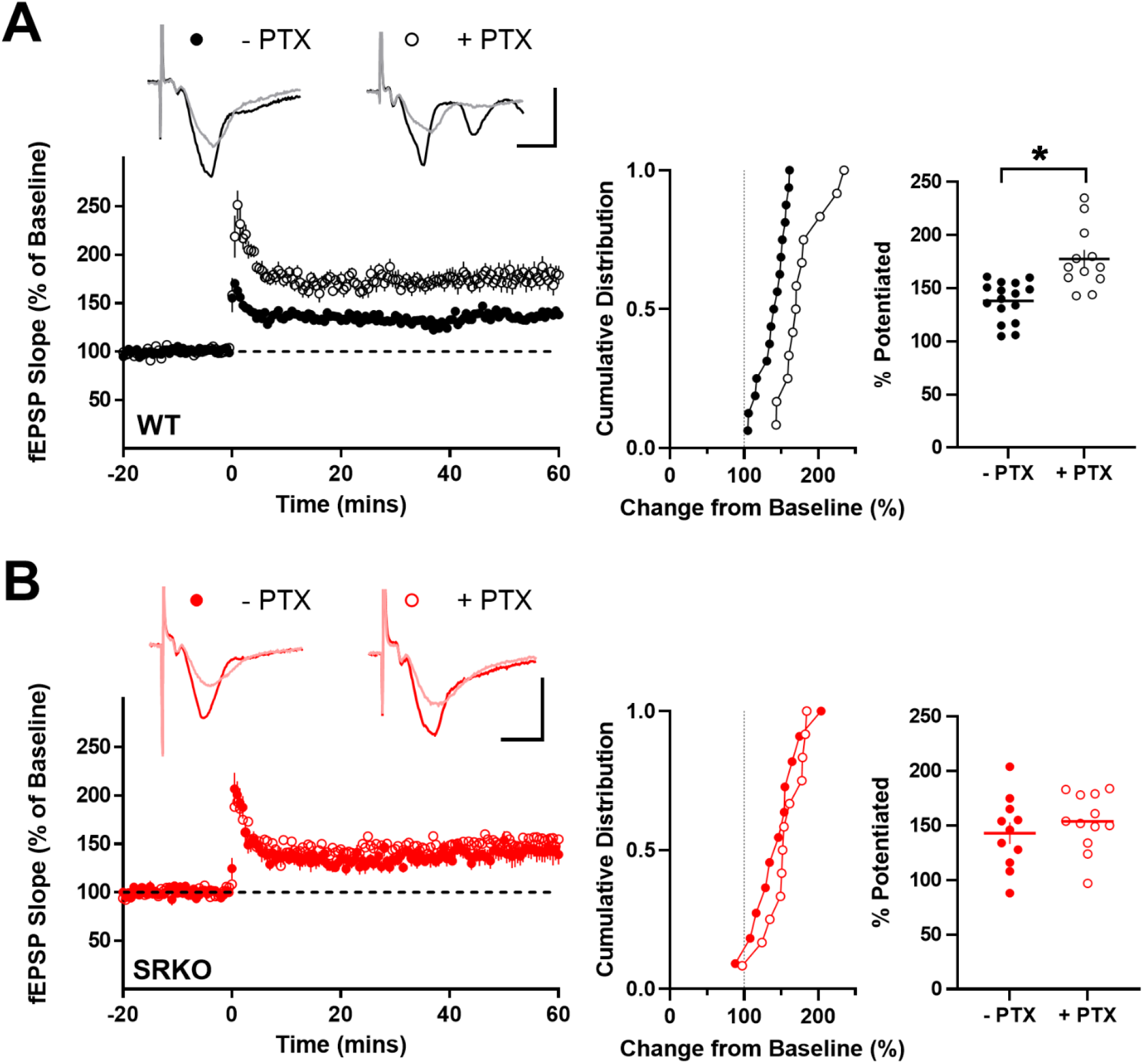
Loss of picrotoxin-induced enhancement of LTP in SRKO mice **(A)** Traces represent superimposed fEPSPs recorded from WT slices during baseline and 60 min after HFS in presence (+) and absence (-) of 50 μM PTX; scale bars: 1 mV, 20 msec. In slices from WT mice, PTX enhances LTP. Middle, cumulative distribution of experiments. Right, summary graph of average percentage potentiation relative to baseline demonstrating that PTX results in significantly enhanced LTP (-PTX: 138±5% of baseline, n=16; +PTX: 178±7% of baseline, n=12; p=0.0002). **(B)** Traces represent superimposed fEPSPs recorded from SRKO slices during baseline and 60 min after HFS in presence (+) and absence (-) of 50 μM PTX; scale bars: 1 mV, 20 msec. In slices from SRKO mice, PTX does not enhance LTP. Middle, cumulative distribution of experiments. Right, summary graph of average percentage potentiation relative to baseline showing no effect of PTX on LTP in slices from SRKO mice (-PTX: 143±10% of baseline, n=11; +PTX: 154±8% of baseline, n=11; p=0.394). Data represent mean ± SEM.

### Reduced inhibitory synapses onto CA1 pyramidal neurons of SRKO mice

To examine the source of the reduced GABAergic inhibition in the SRKO mice, we recorded spontaneous IPSCs (sIPSC) from CA1 pyramidal cells **(Fig. 4A-C)**. There were no significant differences in sIPSC amplitude between SRKO and WT mice (**Fig. 4A**, p=0.138, unpaired t-test, t(34)=1.42), though sIPSC frequency was significantly reduced (**Fig. 4B**, p=0.006, unpaired t-test, t(34)=2.96). Similarly, mIPSC **(Fig. 4D-F)** frequency was significantly reduced in CA1 pyramidal cells from the SRKO mice compared to WT (**Fig. 4E**, p=0.0003, unpaired t-test, t(23)=4.29). There was also a small decrease in mIPSC amplitude in the SRKO neurons (**Fig. 4D**, p=0.042, unpaired t-test, t(23)=2.15). These results suggest that there is a significant reduction of inhibitory synapses onto CA1 pyramidal neurons in the SRKO mice. Though there were no apparent differences in the I/O of excitatory responses at SC-CA1 synapses **(Fig. 1A)**, evoked and spontaneous neurotransmission may be distinct (Kavalali, 2015). Thus, we also examined sEPSCs and mEPSCs from CA1 pyramidal neurons **(Fig. 5)**. We found no significant differences between cells from WT and SRKO mice in sEPSC amplitude (**Fig. 5A**, p=0.79, unpaired t-test, t(22)=0.259), sEPSC frequency (**Fig. 5B**, p=0.47, unpaired t-test, t(22)=0.732), or mEPSC frequency (**Fig. 5D**, p=0.70, unpaired t-test, t(26)=0.383). There was a small, significant increase in mEPSC amplitude in the SRKO cells (**Fig. 5E**, p=0.016, unpaired t-test, t(26)=2.57), that appeared to be most at larger amplitude synapses. Overall, these results, combined with **Figure 1,** suggest that fast excitatory neurotransmission is largely normal in CA1 pyramidal cells from the SRKO mice.

**Figure 4:**
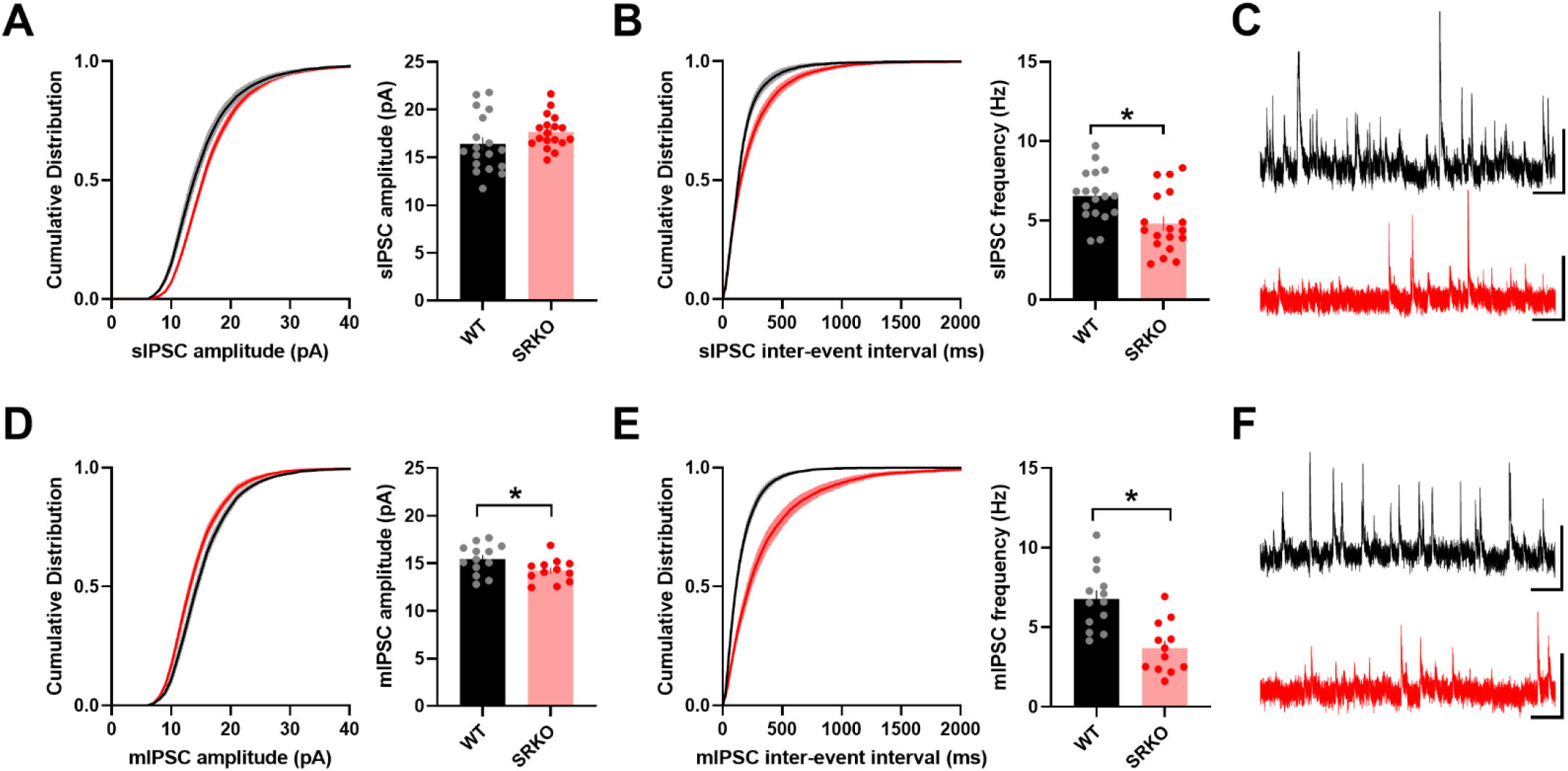
Reduced spontaneous GABAergic synaptic transmission in SRKO mice **(A-C)** Spontaneous IPSCs from CA1 pyramidal cells. **(A)** Cumulative distribution and average amplitude of sIPSCs are unchanged between slices from WT and SRKO mice (WT: 16.41±0.708, n=18; SRKO: 17.66±0.414, n=18; p=0.138). **(B)** The cumulative probability of inter-event intervals reveals a shift towards longer intervals, indicating a decreased frequency of sIPSCs in the SRKO, and the average frequency of sIPSCs was significantly decreased in SRKO compared to WT cells (WT: 6.55±0.38 Hz, n=19; SRKO: 4.80±0.45 Hz, n=18; p=0.006). **(C)** Sample sIPSC traces from WT (black) and SRKO (red) mice; scale bars: 25 pA and 0.5 sec. **(D-F)** Miniature IPSCs from CA1 pyramidal cells **(D)** Cumulative probability and average amplitude of mIPSC were significantly changed between SRKO and WT mice (WT: 15.45 ±0.43 pA, n=13; SRKO: 14.23±0.36 pA, n=12; p<0.042). **(E)** The cumulative probability of inter-event intervals and the average frequency of mIPSCs are significantly decreased in SRKO compared to WT cells (WT: 6.79±0.54 Hz, n=13; SRKO: 3.69±0.47Hz, n=12; p = 0.0003). **(F)** Sample mIPSC traces from WT (black) and SRKO (red) mice; scale bars: 25 pA and 0.5 sec. Data represent mean ± SEM.

**Figure 5:**
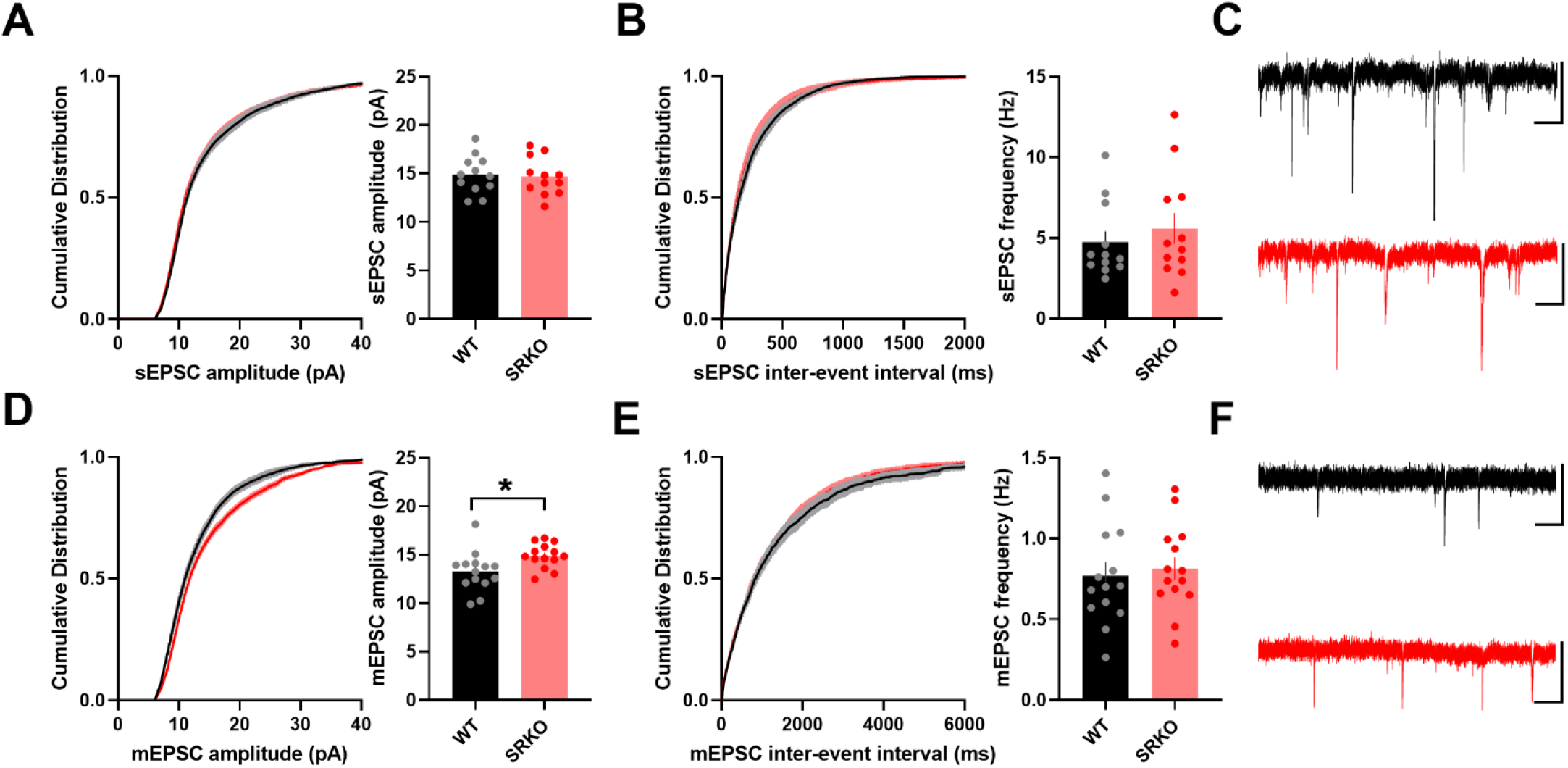
Normal spontaneous excitatory synaptic transmission in SRKO mice. **(A-C)** Spontaneous EPSCs from CA1 pyramidal cells. **(A)** Cumulative probabilityand average amplitudes of sEPSCs were not significantly different between SRKO and WT mice (WT: 14.88±0.56 pA, n=12; SRKO: 14.68±0.56 pA, n=12; p=0.798). **(B)** The cumulative probability of inter-event intervals and average frequency of sEPSCs was also unchanged (WT: 4.74±0.68 Hz, n=12; SRKO: 5.59±0.95 Hz, n=12; p=0.472). **(C)** Sample sEPSC traces from WT (black) and SRKO (red) mice; scale bars: 25 pA and 0.5 sec. **(D-F)** Miniature EPSCs from CA1 pyramidal cells. **(D)** Cumulative probability and average amplitude of mEPSCs were significantly changed between SRKO and WT mice (WT: 13.24±0.54 pA, n=14; SRKO: 14.89±0.34 pA, n=14; p=0.016). **(E)** The cumulative probability of inter-event intervals and average frequency of mEPSCs were not significantly different between SRKO and WT mice (WT: 0.770±0.084 Hz, n=14; SRKO: 0.812±0.072 Hz, n=14; p=0.705). **(F)** Sample mEPSC traces from WT (black) and SRKO (red) mice; scale bars: 25 pA and 0.5 sec. Data represent mean ± SEM.

The reduced frequency of mIPSCs **(Fig. 4E)**, in the absence of apparent changes in presynaptic release probability **(Fig. 1D)**, suggests a reduction in the number of GABAergic synapses onto CA1 pyramidal neurons in the SRKO mice. We then confirmed this synaptic reduction using immunohistochemistry **(Fig. 6)** by staining for the vesicular GABA transporter (VGAT) in hippocampal slices. Consistent with a reduction of synapses from PV+ interneurons, which form perisomatic synapses onto CA1 pyramidal cells, there was a significant reduction of VGAT density (**Fig. 6A-B**, left, p=0.028, unpaired t-test, t(4)=3.36) and intensity (**Fig. 6A-B**, right, p=0.042, unpaired t-test, t(4)=2.95) in the CA1 pyramidal cell layer in the SRKO mice compared with WT. Similarly, in the *stratum radiatum,* there was a nonsignificant reduction in VGAT density (**Fig. 6C-D**, left, p=0.092, unpaired t-test, t(4)=2.21) and a significant decrease in VGAT intensity (**Fig. 6C-D**, right, p=0.024, unpaired t-test, t(4)=3.53), that was evenly distributed throughout the *stratum radiatum* **(Fig. 6E)** suggesting a broader GABAergic synapse deficit. Taken together with the significant reduction in mIPSC frequency, these results suggest that a loss of GABAergic synapse density in the hippocampus underlies the increased E/I ratio in the SRKO mice.

**Figure 6:**
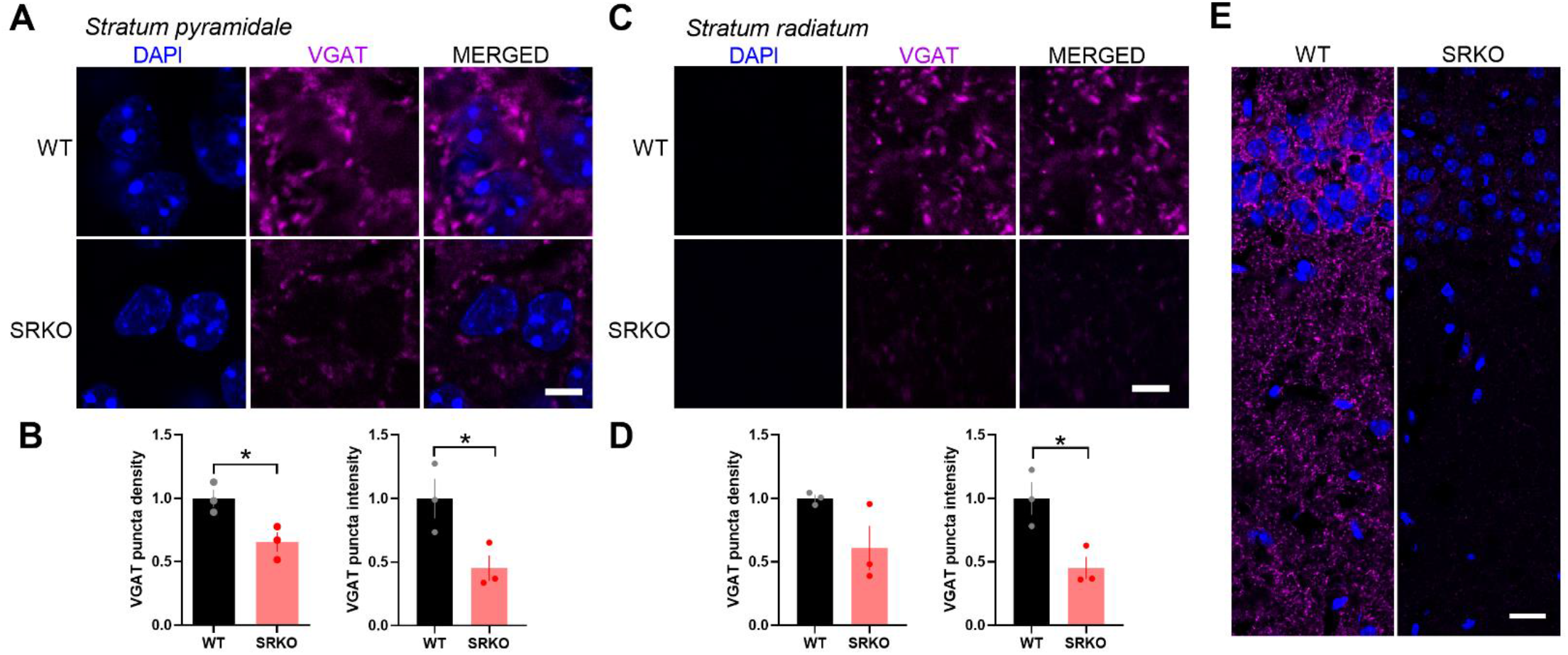
Reduced GABAergic synapses onto CA1 pyramidal neurons in SRKO mice. **(A**) Representative images of VGAT labeling in the *stratum pyramidale* of CA1 hippocampus show a reduction in VGAT antibody labeling in SRKO mice; scale bar indicates 5 μm. **(B)** Both normalized average VGAT puncta density (WT: 1.000±0.069, n=3; SRKO: 0.654±0.076, n=3; p=0.028) and normalized average VGAT puncta intensity in CA1 *stratum pyramidale* (WT: 1.000±0.156, n=3; SRKO: 0.453±0.101, n=3; p=0.042) are significantly lower in SRKO mice. **(C)** Representative images of VGAT labeling in the *stratum radiatum* of CA1 hippocampus show a reduction in VGAT labeling in SRKO mice; scale bar indicates 5 μm. **(D)** There is a non-significant reduction in the normalized average VGAT puncta density in *stratum radiatum* of CA1 of SRKO mice (WT: 1.000±0.028, n=3; SRKO: 0.609±0.175, n=3; p=0.092), while the normalized average VGAT puncta intensity in *stratum radiatum* is significantly reduced in the SRKO mice (WT: 1.000±0.128, n=3; SRKO: 0.452±0.088, n=3; p=0.024). **(E)** Representative images of hippocampal CA1 show that the reduction in VGAT signal is consistent across strata of CA1 in SRKO mice; scale bar indicates 20 μm. Data represent mean ± SEM.

### Deletion of SR from CA1 pyramidal neurons results in a cell-autonomous reduction in GABAergic synapses

Early studies suggested that D-serine is exclusively synthesized and released by astrocytes (Schell et al., 1995; Schell et al., 1997; Wolosker et al., 1999) leading to the classification of D-serine as a gliotransmitter (Wolosker et al., 2002; Miller, 2004; Panatier et al., 2006). More recent studies, using the SR knockout mice as controls, have strongly supported a predominantly neuronal localization (Kartvelishvily et al., 2006; Yoshikawa et al., 2007; Miya et al., 2008; Basu et al., 2009; Ding et al., 2011; Ehmsen et al., 2013; Balu et al., 2014; Wolosker et al., 2016; Balu et al., 2018). Furthermore, in agreement with previous studies in cultured neurons (Ma et al., 2014; Lin et al., 2016), we recently reported that SR localizes to the apical dendrites and the post-synaptic density *in situ* in hippocampal CA1 pyramidal neurons and regulates postsynaptic NMDARs (Wong et al., 2020). Importantly, while conditional knockout (cKO) of SR from astrocytes has minimal impact on SR levels, cKO from CaMKIIα-expressing forebrain glutamatergic neurons results in ~65% reduction of SR expression in the cortex and hippocampus (Benneyworth et al., 2012). The remainder of SR expression is thought to be from GABAergic interneurons. As such, we sought to determine if the decrease in GABAergic synapses onto CA1 pyramidal neurons in the SRKO mice was due to the loss of SR in the pyramidal cells themselves. We utilized a single-neuron genetic approach in the SR^fl^ mice in which SR was removed in a sparse subset of CA1 pyramidal neurons by neonatal stereotaxic injection of adeno-associated virus, serotype 1 expressing a Cre recombinase GFP fusion protein (AAV1-Cre:GFP) **(Fig. 7A)**. This mosaic transduction allows for whole-cell recordings from Cre-expressing (Cre) and untransduced neurons (Ctrl) **(Fig. 7B)** providing a measurement of the cell-autonomous effects of SR deletion. Similar to the SRKO mice **(Fig. 4)**, we found no differences in mIPSC amplitude (**Fig. 7C**, p=0.939, unpaired t-test, t(19)=2.022), but significantly reduced mIPSC frequency (**Fig. 7D**, p=0.039, unpaired t-test, t(19)=2.218) in Cre-expressing CA1 pyramidal neurons compared to control neurons. These results suggest that cKO of SR from CA1 pyramidal neurons results in a cell-autonomous reduction in GABAergic synapses.

**Figure 7:**
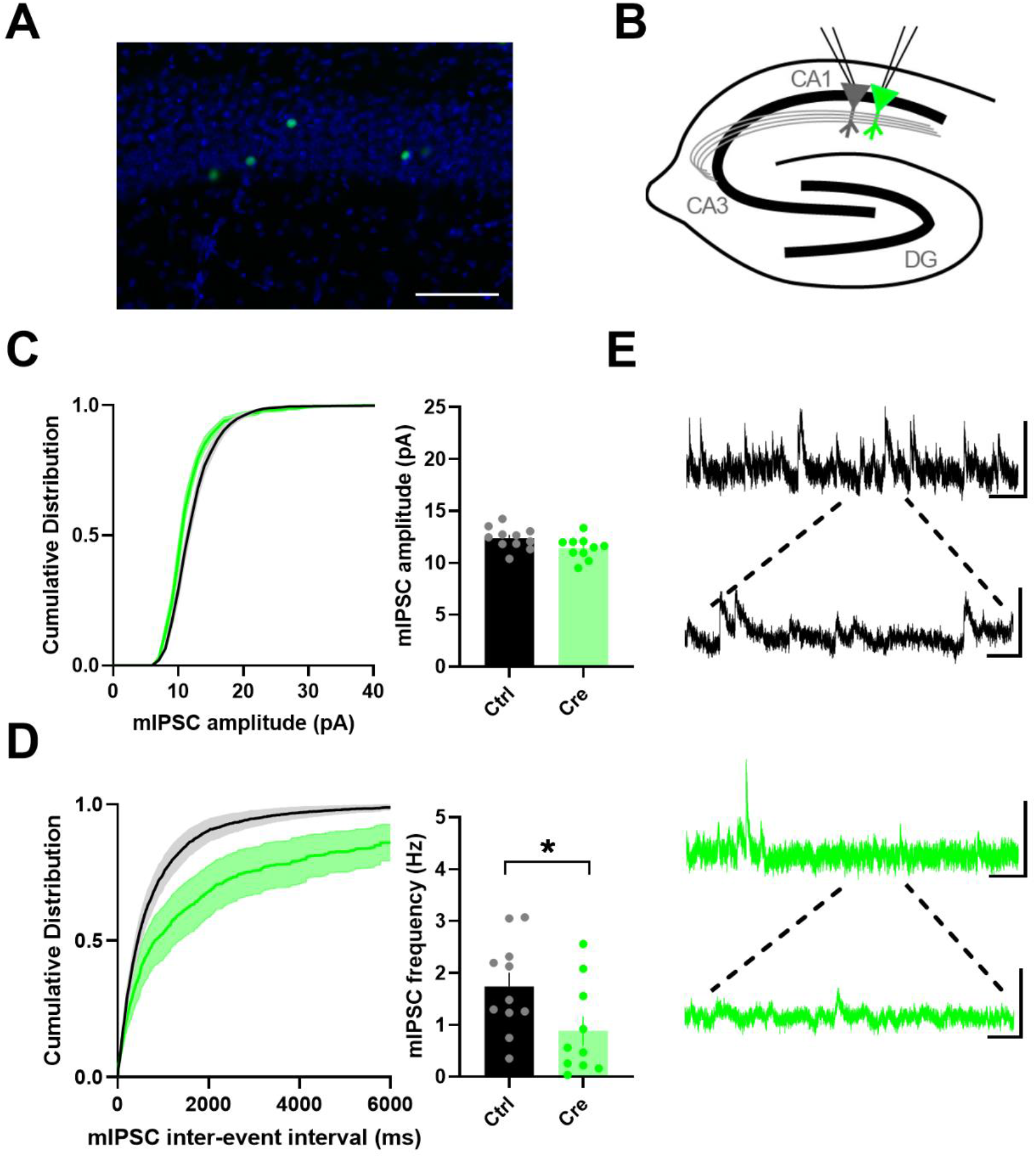
Cell-autonomous reductions in spontaneous GABAergic synaptic transmission onto CA1 pyramidal cells following single-neuron SR deletion **(A)** Representative image of the sparse transduction of CA1 pyramidal cells by AAV1-Cre:GFP counterstained by DAPI. Scale bar indicates 100 μm. **(B)** Schematic of experimental setup. Simultaneous whole-cell mIPSC recordings from were made from neighboring transduced (Cre+) and control CA1 pyramidal cells. **(C)** Cumulative probability and average mIPSC amplitude from Cre+ neurons was not significantly different than those from control cells (WT: 12.37±0.32 pA, n=11; SRKO: 11.42±0.34 pA, n=10; p=0.939). **(D)** Cumulative probability of inter-event intervals and average frequency of mIPSCs from Cre+ neurons was significantly decreased compared to control cells (WT: 1.74±0.27Hz, n=11; SRKO: 0.88±0.28 Hz, n=10; p=0.039. **(E)** Sample mIPSC traces from control (black, top) and Cre+ (green, bottom) pyramidal neurons; scale bars: 25 pA and 0.5 sec; inset, 25 pA and 100 ms. Data represent mean ± SEM.

## Discussion

Broad NMDAR deletion causes overly pronounced phenotypes that do not adequately model schizophrenia (Nakazawa et al., 2017). Germline deletion of NMDARs from mice is perinatally lethal (Forrest et al., 1994; Li et al., 1994; Kutsuwada et al., 1996) and embryonic deletion from only forebrain pyramidal neurons results in death within the first month (Iwasato et al., 2000; Ultanir et al., 2007; Quintero et al., 2008). Similarly, mice with a homozygous embryonic deletion of NMDARs from migrating forebrain GABAergic neurons expressing the Dlx5/6 promoter (Zerucha et al., 2000), are reportedly nonviable (Nakazawa et al., 2017). Moreover, broad and regional deletion of NMDARs severely disrupts cortical patterning during development (Li et al., 1994; Iwasato et al., 2000). The NMDAR hypomorph mice (Mohn et al., 1999), which have only 5-10% of wildtype NMDAR expression, have been hailed as a major transgenic model of the NMDAR hypofunction in schizophrenia (Gainetdinov et al., 2001), though they have also been highly criticized for having more global cognitive impairments with earlier onset than what is seen in schizophrenia (Barkus et al., 2012; Gandal et al., 2012; Moy et al., 2012). Interestingly, decreases in NMDAR protein is not a consistent finding in schizophrenia (Catts et al., 2016), suggesting that the hypofunction may be more functional (e.g. downstream signaling) than structural (Banerjee et al., 2015). Indeed, NMDARs are macromolecular machines (Fan et al., 2014) involved in a plethora of signaling processes in neurons and complete loss of NMDARs could lead to a broad range of allostatic changes. In this study, we utilized a mouse model of NMDAR hypofunction that involves a functional rather than structural reduction in NMDAR activity, the SRKO mice (Basu et al., 2009). In the SRKO mice, there is a >90% decrease in the levels of D-serine, the primary co-agonist for synaptic NMDARs in the forebrain (Mothet et al., 2000; Basu et al., 2009). Indeed, deficiency of D-serine and the subsequent hypofunction of NMDARs has been implicated in the pathophysiology of schizophrenia (Coyle, 2012) and the SRKO mice display many well-characterized hallmarks of schizophrenia, including reductions in dendritic complexity and spine density (Rosoklija et al., 2000; Balu et al., 2012; Balu et al., 2013) and impaired performance on various cognitive tasks (Basu et al., 2009; Balu et al., 2013).

Using the SRKO mice, we have explored the relationship between NMDAR hypofunction and GABAergic inhibition. Because interneurons expressing the calcium-binding protein parvalbumin (PV+) are particularly affected in schizophrenia (Hashimoto et al., 2003; Hashimoto et al., 2008; Mellios et al., 2009), previous studies have examined PV expression in the SRKO mice. While one study reported a 26% reduction in PV+ cells in the anterior cingulate cortex of the SRKO mice (Steullet et al., 2017), another found no change in PV immunoreactivity in the hippocampus, prelimbic and infralimbic cortices (Benneyworth et al., 2011). However, using electrophysiological approaches in *ex vivo* hippocampal slices we found a significant reduction of GABAergic synapses onto CA1 pyramidal neurons in the SRKO mice. This reduction of GABAergic synaptic inhibition onto pyramidal cells increases the E/I balance resulting in enhanced synaptically-driven neuronal excitability and elevated broad-spectrum oscillatory activity *in vitro*. These results are consistent with a recent study demonstrating disruptions in the auditory steady-state response in the SRKO mice (Balla et al., 2020), which suggests impaired coordination between pyramidal cells and interneurons.

We further show, using a single-neuron genetic deletion approach, that the loss of GABAergic synapses onto pyramidal neurons observed in the SRKO mice is driven in a cell-autonomous manner following the deletion of SR in individual CA1 pyramidal cells. Indeed, recent studies have shown a critical role for NMDARs on pyramidal neurons in regulating GABAergic synapse development (Lu et al., 2013; Gu et al., 2016; Gu and Lu, 2018).

Specifically, deletion of the obligatory GluN1 subunit of NMDARs from single CA1 pyramidal cells in early development leads to a significant reduction in mIPSC frequency and a loss of GABAergic synapses (Gu et al., 2016). Importantly, a similar loss of GABAergic synapses upon GluN1 deletion was observed in layer 2/3 pyramidal neurons in the motor cortex and midbrain dopaminergic neurons in the ventral tegmental area (Gu and Lu, 2018), suggesting a more generalizable mechanism. This work builds upon older pharmacological studies showing that NMDAR activity can accelerate GABAergic synapse development (Harris et al., 1995; Aamodt et al., 2000; Henneberger et al., 2005; Lin et al., 2008). Interestingly, NMDARs have been found to co-localize with GABAA receptors at GABAergic synapses in the developing brain (Gundersen et al., 2004; Szabadits et al., 2011; Cserep et al., 2012), though the function of this localization remains unclear. Together, these results support a model whereby NMDAR hypofunction on pyramidal neurons can lead to GABAergic dysfunction through a loss of GABAergic synapses.

The cellular location of the NMDAR hypofunction in schizophrenia has been intensely studied yet remains poorly understood. A large body of pharmacological studies using uncompetitive NMDAR antagonists support a locus of NMDAR hypofunction on cortical GABAergic interneurons, particularly PV positive cells (Hashimoto et al., 2003; Hashimoto et al., 2008; Mellios et al., 2009). Notably, acute systemic administration of NMDAR antagonists results in the increased activity of cortical pyramidal neurons (Suzuki et al., 2002; Jackson et al., 2004), spillover of cortical glutamate (Moghaddam et al., 1997; Lorrain et al., 2003), and increases in cortical gamma power (Driesen et al., 2013; Hunt and Kasicki, 2013), indicative of increased E/I balance and pyramidal cell disinhibition. Similar evidence for increased cortical excitability following administration of NMDAR antagonists have been found in human studies (Lahti et al., 1995a; Lahti et al., 1995b; Breier et al., 1997; Vollenweider et al., 1997). These findings are consistent with the increase in E/I balance and disinhibition we observe here in the SRKO mice; however, NMDAR antagonists are thought to preferentially inhibit receptors on fast-spiking PV-positive interneurons (Homayoun and Moghaddam, 2007).

Cell-type-specific knockouts of GluN1 from either pyramidal neurons or PV+ interneurons have provided additional insights into the locus of NMDAR hypofunction in schizophrenia. For example, deletion of GluN1 from PV+ interneurons leads to cortical and hippocampal disinhibition and an increase in the baseline gamma power in the hippocampus (Korotkova et al., 2010; Carlen et al., 2012). In addition, acute MK801-induced behaviors were not detected in these mice (Carlen et al., 2012), providing decisive evidence for PV+ interneurons being the locus of NMDAR hypofunction upon systemic NMDAR antagonist administration in adult rodents. Behaviorally, these mice have selective impairments in working memory, habituation, and sociability, but display normal pre-pulse inhibition (PPI) (Korotkova et al., 2010; Carlen et al., 2012; Saunders et al., 2013). Importantly, because PV promoter expression, and thus NMDAR removal, begins at 2-4 weeks of age (Taniguchi et al., 2011; Carlen et al., 2012; Saunders et al., 2013), these mice may not fully model the neurodevelopmental changes occurring in schizophrenia.

Similarly, mice with a deletion of GluN1 from forebrain pyramidal neurons using the CaMKII promoter display a variety of schizophrenia-related phenotypes, including reductions in social interaction, nest-building, and spatial working memory (McHugh et al., 1996; Tatard-Leitman et al., 2015). Interestingly, there was also an increase in locomotor activity in the CaMKII-Cre/GluN1 KO mice consistent with dopaminergic models of psychosis (van den Buuse, 2010; Tatard-Leitman et al., 2015). Similar to our results, CA1 pyramidal cell excitability was increased along with increased broadband local field potential power in the CaMKII-Cre/GluN1 KO mice (Tatard-Leitman et al., 2015); however, this was an increase in intrinsic excitability attributable to a reduction in GIRK2 channel activity, rather than due to the loss of synaptic inhibition seen here. Furthermore, no changes in mRNA levels were found in the hippocampus for the GABAergic markers GAD67, PV, cholecystokinin, and somatostatin (Tatard-Leitman et al., 2015), suggesting a lack of effects on inhibition. Importantly, the CaMKII promoter drives GluN1 deletion in these mice beginning at 3-4 weeks of age in CA1 pyramidal neurons which then spreads more broadly throughout the forebrain by 4 months (Tsien et al., 1996). Thus, as with the deletion of GluN1 from PV+ interneurons, these mice may not recapitulate the developmental aspects of NMDAR hypofunction.

The chronic, developmental loss of inhibitory synapses onto hippocampal pyramidal cells could lead to the GABAergic and oscillatory dysfunction seen in schizophrenia (Gonzalez-Burgos and Lewis, 2012). In the Pyramidal-Interneuron Network Gamma (PING) circuit model for gamma oscillations, phasic excitatory input from pyramidal cells is the main source of PV+ interneuron activation and the feedback inhibition of the pyramidal neuron leads to network synchrony (Tiesinga and Sejnowski, 2009). Indeed, modeling has shown that the shunting inhibition by GABAergic synapses, especially perisomatic synapses and those along the proximal apical dendrite, promotes neuronal synchronization (Bartos et al., 2007). Thus, disruption of these feedback synapses onto pyramidal cells could result in a combination of increased excitation, as seen in broad-band baseline power increases along with disrupted task-evoked synchrony.

Consistent with a reduction in synapses from PV+ basket cells, we found a significant reduction in perisomatic VGAT puncta density and intensity in the CA1 pyramidal cell layer. However, the density and intensity of VGAT puncta were also decreased in the stratum radiatum with no apparent proximal-distal differences along the apical dendrites of CA1 pyramidal neurons, supporting a broad reduction of GABAergic synapses. Indeed, while PV+ interneurons are particularly affected in schizophrenia (Hashimoto et al., 2003; Hashimoto et al., 2008; Mellios et al., 2009), multiple interneuron subtypes have been implicated (Benes et al., 2008; Hashimoto et al., 2008; Morris et al., 2008; Beneyto et al., 2012) and hippocampal inhibitory networks appear especially sensitive to NMDAR hypofunction (Ling and Benardo, 1995; Grunze et al., 1996). Overall, our data suggest that a pyramidal cell locus of synaptic NMDAR hypofunction could lead to GABAergic deficits through the impaired development of feedback inhibitory synapses. Additional studies will be needed to elucidate the molecular mechanisms underlying the role of NMDARs in GABAergic synapse development and to ascertain the relationship between inhibitory synapses on pyramidal neurons and endophenotypes in schizophrenia.

## Acknowledgements

This work was supported by R21MH116315 and R01MH117130 to JAG. We would like to thank Joseph Coyle for the SR floxed and knockout mice and Haley Martin, Zaiyang “Sunny” Zhang, and Casey Martin for their assistance in mouse breeding and genotyping.

## References

Aamodt SM, Shi J, Colonnese MT, Veras W, Constantine-Paton M (2000) Chronic NMDA exposure accelerates development of GABAergic inhibition in the superior colliculus. J Neurophysiol 83:1580–1591.

Balla A, Ginsberg SD, Abbas AI, Sershen H, Javitt DC (2020) Translational neurophysiological biomarkers of N-methyl-d-aspartate receptor dysfunction in serine racemase knockout mice. Biomarkers in Neuropsychiatry 2:100019.

Balu DT, Coyle JT (2014) Chronic D-serine reverses arc expression and partially rescues dendritic abnormalities in a mouse model of NMDA receptor hypofunction. Neurochem Int 75:76–78.

Balu DT, Basu AC, Corradi JP, Cacace AM, Coyle JT (2012) The NMDA receptor co-agonists, D-serine and glycine, regulate neuronal dendritic architecture in the somatosensory cortex. Neurobiol Dis 45:671–682.

Balu DT, Takagi S, Puhl MD, Benneyworth MA, Coyle JT (2014) D-serine and serine racemase are localized to neurons in the adult mouse and human forebrain. Cell Mol Neurobiol 34:419–435.

Balu DT, Li Y, Puhl MD, Benneyworth MA, Basu AC, Takagi S, Bolshakov VY, Coyle JT (2013) Multiple risk pathways for schizophrenia converge in serine racemase knockout mice, a mouse model of NMDA receptor hypofunction. Proc Natl Acad Sci U S A 110:E2400–2409.

Balu DT, Presti KT, Huang CCY, Muszynski K, Radzishevsky I, Wolosker H, Guffanti G, Ressler KJ, Coyle JT (2018) Serine Racemase and D-serine in the Amygdala Are Dynamically Involved in Fear Learning. Biol Psychiatry 83:273–283.

Balu DT, Li Y, Takagi S, Presti KT, Ramikie TS, Rook JM, Jones CK, Lindsley CW, Conn PJ, Bolshakov VY, Coyle JT (2016) An mGlu5-Positive Allosteric Modulator Rescues the Neuroplasticity Deficits in a Genetic Model of NMDA Receptor Hypofunction in Schizophrenia. Neuropsychopharmacology 41:2052–2061.

Banerjee A, Wang HY, Borgmann-Winter KE, MacDonald ML, Kaprielian H, Stucky A, Kvasic J, Egbujo C, Ray R, Talbot K, Hemby SE, Siegel SJ, Arnold SE, Sleiman P, Chang X, Hakonarson H, Gur RE, Hahn CG (2015) Src kinase as a mediator of convergent molecular abnormalities leading to NMDAR hypoactivity in schizophrenia. Mol Psychiatry 20:1091–1100.

Barkus C, Dawson LA, Sharp T, Bannerman DM (2012) GluN1 hypomorph mice exhibit wide-ranging behavioral alterations. Genes Brain Behav 11:342–351.

Bartley AF, Lucas EK, Brady LJ, Li Q, Hablitz JJ, Cowell RM, Dobrunz LE (2015) Interneuron Transcriptional Dysregulation Causes Frequency-Dependent Alterations in the Balance of Inhibition and Excitation in Hippocampus. J Neurosci 35:15276–15290.

Bartos M, Vida I, Jonas P (2007) Synaptic mechanisms of synchronized gamma oscillations in inhibitory interneuron networks. Nat Rev Neurosci 8:45–56.

Basu AC, Tsai GE, Ma CL, Ehmsen JT, Mustafa AK, Han L, Jiang ZI, Benneyworth MA, Froimowitz MP, Lange N, Snyder SH, Bergeron R, Coyle JT (2009) Targeted disruption of serine racemase affects glutamatergic neurotransmission and behavior. Mol Psychiatry 14:719–727.

Bateup HS, Johnson CA, Denefrio CL, Saulnier JL, Kornacker K, Sabatini BL (2013) Excitatory/inhibitory synaptic imbalance leads to hippocampal hyperexcitability in mouse models of tuberous sclerosis. Neuron 78:510–522.

Bendikov I, Nadri C, Amar S, Panizzutti R, De Miranda J, Wolosker H, Agam G (2007) A CSF and postmortem brain study of D-serine metabolic parameters in schizophrenia. Schizophr Res 90:41–51.

Benes FM, Lim B, Matzilevich D, Subburaju S, Walsh JP (2008) Circuitry-based gene expression profiles in GABA cells of the trisynaptic pathway in schizophrenics versus bipolars. Proc Natl Acad Sci U S A 105:20935–20940.

Beneyto M, Morris HM, Rovensky KC, Lewis DA (2012) Lamina- and cell-specific alterations in cortical somatostatin receptor 2 mRNA expression in schizophrenia. Neuropharmacology 62:1598–1605.

Benneyworth MA, Roseman AS, Basu AC, Coyle JT (2011) Failure of NMDA receptor hypofunction to induce a pathological reduction in PV-positive GABAergic cell markers. Neurosci Lett 488:267–271.

Benneyworth MA, Li Y, Basu AC, Bolshakov VY, Coyle JT (2012) Cell selective conditional null mutations of serine racemase demonstrate a predominate localization in cortical glutamatergic neurons. Cell Mol Neurobiol 32:613–624.

Bickel S, Lipp HP, Umbricht D (2008) Early auditory sensory processing deficits in mouse mutants with reduced NMDA receptor function. Neuropsychopharmacology 33:1680–1689.

Bischofberger J, Engel D, Li L, Geiger JR, Jonas P (2006) Patch-clamp recording from mossy fiber terminals in hippocampal slices. Nat Protoc 1:2075–2081.

Breier A, Malhotra AK, Pinals DA, Weisenfeld NI, Pickar D (1997) Association of ketamine-induced psychosis with focal activation of the prefrontal cortex in healthy volunteers. Am J Psychiatry 154:805–811.

Carlen M, Meletis K, Siegle JH, Cardin JA, Futai K, Vierling-Claassen D, Ruhlmann C, Jones SR, Deisseroth K, Sheng M, Moore CI, Tsai LH (2012) A critical role for NMDA receptors in parvalbumin interneurons for gamma rhythm induction and behavior. Mol Psychiatry 17:537–548.

Catts VS, Lai YL, Weickert CS, Weickert TW, Catts SV (2016) A quantitative review of the postmortem evidence for decreased cortical N-methyl-D-aspartate receptor expression levels in schizophrenia: How can we link molecular abnormalities to mismatch negativity deficits? Biol Psychol 116:57–67.

Cho KK, Hoch R, Lee AT, Patel T, Rubenstein JL, Sohal VS (2015) Gamma rhythms link prefrontal interneuron dysfunction with cognitive inflexibility in Dlx5/6(+/-) mice. Neuron 85:1332–1343.

Chumakov I et al. (2002) Genetic and physiological data implicating the new human gene G72 and the gene for D-amino acid oxidase in schizophrenia. Proc Natl Acad Sci U S A 99:13675–13680.

Coyle JT (2012) NMDA receptor and schizophrenia: a brief history. Schizophr Bull 38:920–926.

Coyle JT, Balu DT (2018) The Role of Serine Racemase in the Pathophysiology of Brain Disorders. Adv Pharmacol 82:35–56.

Cserep C, Szabadits E, Szonyi A, Watanabe M, Freund TF, Nyiri G (2012) NMDA receptors in GABAergic synapses during postnatal development. PLoS One 7:e37753.

Detera-Wadleigh SD, McMahon FJ (2006) G72/G30 in schizophrenia and bipolar disorder: review and meta-analysis. Biol Psychiatry 60:106–114.

DeVito LM, Balu DT, Kanter BR, Lykken C, Basu AC, Coyle JT, Eichenbaum H (2011) Serine racemase deletion disrupts memory for order and alters cortical dendritic morphology. Genes Brain Behav 10:210–222.

Ding X, Ma N, Nagahama M, Yamada K, Semba R (2011) Localization of D-serine and serine racemase in neurons and neuroglias in mouse brain. Neurol Sci 32:263–267.

Driesen NR, McCarthy G, Bhagwagar Z, Bloch M, Calhoun V, D’Souza DC, Gueorguieva R, He G, Ramachandran R, Suckow RF, Anticevic A, Morgan PT, Krystal JH (2013) Relationship of resting brain hyperconnectivity and schizophrenia-like symptoms produced by the NMDA receptor antagonist ketamine in humans. Mol Psychiatry 18:1199–1204.

Duncan G, Miyamoto S, Gu H, Lieberman J, Koller B, Snouwaert J (2002) Alterations in regional brain metabolism in genetic and pharmacological models of reduced NMDA receptor function. Brain Res 951:166–176.

Duncan GE, Moy SS, Lieberman JA, Koller BH (2006) Typical and atypical antipsychotic drug effects on locomotor hyperactivity and deficits in sensorimotor gating in a genetic model of NMDA receptor hypofunction. Pharmacol Biochem Behav 85:481–491.

Duncan GE, Moy SS, Perez A, Eddy DM, Zinzow WM, Lieberman JA, Snouwaert JN, Koller BH (2004) Deficits in sensorimotor gating and tests of social behavior in a genetic model of reduced NMDA receptor function. Behav Brain Res 153:507–519.

Dzirasa K, Ramsey AJ, Takahashi DY, Stapleton J, Potes JM, Williams JK, Gainetdinov RR, Sameshima K, Caron MG, Nicolelis MA (2009) Hyperdopaminergia and NMDA receptor hypofunction disrupt neural phase signaling. J Neurosci 29:8215–8224.

Ehmsen JT, Ma TM, Sason H, Rosenberg D, Ogo T, Furuya S, Snyder SH, Wolosker H (2013) D-serine in glia and neurons derives from 3-phosphoglycerate dehydrogenase. J Neurosci 33:12464–12469.

Elmer BM, Estes ML, Barrow SL, McAllister AK (2013) MHCI requires MEF2 transcription factors to negatively regulate synapse density during development and in disease. J Neurosci 33:13791–13804.

Fan X, Jin WY, Wang YT (2014) The NMDA receptor complex: a multifunctional machine at the glutamatergic synapse. Front Cell Neurosci 8:160.

Fellous JM, Sejnowski TJ (2000) Cholinergic induction of oscillations in the hippocampal slice in the slow (0.5-2 Hz), theta (5-12 Hz), and gamma (35-70 Hz) bands. Hippocampus 10:187–197.

Fisahn A, Pike FG, Buhl EH, Paulsen O (1998) Cholinergic induction of network oscillations at 40 Hz in the hippocampus in vitro. Nature 394:186–189.

Forrest D, Yuzaki M, Soares HD, Ng L, Luk DC, Sheng M, Stewart CL, Morgan JI, Connor JA, Curran T (1994) Targeted disruption of NMDA receptor 1 gene abolishes NMDA response and results in neonatal death. Neuron 13:325–338.

Fradley RL, O’Meara GF, Newman RJ, Andrieux A, Job D, Reynolds DS (2005) STOP knockout and NMDA NR1 hypomorphic mice exhibit deficits in sensorimotor gating. Behav Brain Res 163:257–264.

Gainetdinov RR, Mohn AR, Caron MG (2001) Genetic animal models: focus on schizophrenia. Trends Neurosci 24:527–533.

Gandal MJ, Anderson RL, Billingslea EN, Carlson GC, Roberts TP, Siegel SJ (2012) Mice with reduced NMDA receptor expression: more consistent with autism than schizophrenia? Genes Brain Behav 11:740–750.

Glausier JR, Lewis DA (2017) GABA and schizophrenia: Where we stand and where we need to go. Schizophr Res 181:2–3.

Goltsov AY, Loseva JG, Andreeva TV, Grigorenko AP, Abramova LI, Kaleda VG, Orlova VA, Moliaka YK, Rogaev EI (2006) Polymorphism in the 5’-promoter region of serine racemase gene in schizophrenia. Mol Psychiatry 11:325–326.

Gonzalez-Burgos G, Lewis DA (2012) NMDA receptor hypofunction, parvalbumin-positive neurons, and cortical gamma oscillations in schizophrenia. Schizophr Bull 38:950–957.

Gonzalez-Burgos G, Fish KN, Lewis DA (2011) GABA neuron alterations, cortical circuit dysfunction and cognitive deficits in schizophrenia. Neural Plast 2011:723184.

Gray JA, Shi Y, Usui H, During MJ, Sakimura K, Nicoll RA (2011) Distinct modes of AMPA receptor suppression at developing synapses by GluN2A and GluN2B: single-cell NMDA receptor subunit deletion in vivo. Neuron 71:1085–1101.

Grunze HC, Rainnie DG, Hasselmo ME, Barkai E, Hearn EF, McCarley RW, Greene RW (1996) NMDA-dependent modulation of CA1 local circuit inhibition. J Neurosci 16:2034–2043.

Gu X, Lu W (2018) Genetic deletion of NMDA receptors suppresses GABAergic synaptic transmission in two distinct types of central neurons. Neurosci Lett 668:147–153.

Gu X, Zhou L, Lu W (2016) An NMDA Receptor-Dependent Mechanism Underlies Inhibitory Synapse Development. Cell Rep 14:471–478.

Gulledge AT, Kampa BM, Stuart GJ (2005) Synaptic integration in dendritic trees. J Neurobiol 64:75–90.

Gundersen V, Holten AT, Storm-Mathisen J (2004) GABAergic synapses in hippocampus exocytose aspartate on to NMDA receptors: quantitative immunogold evidence for co-transmission. Mol Cell Neurosci 26:156–165.

Hajos N, Ellender TJ, Zemankovics R, Mann EO, Exley R, Cragg SJ, Freund TF, Paulsen O (2009) Maintaining network activity in submerged hippocampal slices: importance of oxygen supply. Eur J Neurosci 29:319–327.

Halene TB, Ehrlichman RS, Liang Y, Christian EP, Jonak GJ, Gur TL, Blendy JA, Dow HC, Brodkin ES, Schneider F, Gur RC, Siegel SJ (2009) Assessment of NMDA receptor NR1 subunit hypofunction in mice as a model for schizophrenia. Genes Brain Behav 8:661–675.

Harris BT, Costa E, Grayson DR (1995) Exposure of neuronal cultures to K+ depolarization or to N-methyl-D-aspartate increases the transcription of genes encoding the alpha 1 and alpha 5 GABAA receptor subunits. Brain Res Mol Brain Res 28:338–342.

Hashimoto T, Volk DW, Eggan SM, Mirnics K, Pierri JN, Sun Z, Sampson AR, Lewis DA (2003) Gene expression deficits in a subclass of GABA neurons in the prefrontal cortex of subjects with schizophrenia. J Neurosci 23:6315–6326.

Hashimoto T, Arion D, Unger T, Maldonado-Aviles JG, Morris HM, Volk DW, Mirnics K, Lewis DA (2008) Alterations in GABA-related transcriptome in the dorsolateral prefrontal cortex of subjects with schizophrenia. Mol Psychiatry 13:147–161.

Henneberger C, Papouin T, Oliet SH, Rusakov DA (2010) Long-term potentiation depends on release of D-serine from astrocytes. Nature 463:232–236.

Henneberger C, Juttner R, Schmidt SA, Walter J, Meier JC, Rothe T, Grantyn R (2005) GluR- and TrkB-mediated maturation of GABA receptor function during the period of eye opening. Eur J Neurosci 21:431–440.

Heresco-Levy U, Javitt DC, Ebstein R, Vass A, Lichtenberg P, Bar G, Catinari S, Ermilov M (2005) D-serine efficacy as add-on pharmacotherapy to risperidone and olanzapine for treatment-refractory schizophrenia. Biol Psychiatry 57:577–585.

Homayoun H, Moghaddam B (2007) NMDA receptor hypofunction produces opposite effects on prefrontal cortex interneurons and pyramidal neurons. J Neurosci 27:11496–11500.

Hunt MJ, Kasicki S (2013) A systematic review of the effects of NMDA receptor antagonists on oscillatory activity recorded in vivo. J Psychopharmacol 27:972–986.

Iwasato T, Datwani A, Wolf AM, Nishiyama H, Taguchi Y, Tonegawa S, Knopfel T, Erzurumlu RS, Itohara S (2000) Cortex-restricted disruption of NMDAR1 impairs neuronal patterns in the barrel cortex. Nature 406:726–731.

Jackson ME, Homayoun H, Moghaddam B (2004) NMDA receptor hypofunction produces concomitant firing rate potentiation and burst activity reduction in the prefrontal cortex. Proc Natl Acad Sci U S A 101:8467–8472.

Javitt DC, Zukin SR (1991) Recent advances in the phencyclidine model of schizophrenia. Am J Psychiatry 148:1301–1308.

Jentsch JD, Redmond DE, Jr., Elsworth JD, Taylor JR, Youngren KD, Roth RH (1997) Enduring cognitive deficits and cortical dopamine dysfunction in monkeys after long-term administration of phencyclidine. Science 277:953–955.

Kartvelishvily E, Shleper M, Balan L, Dumin E, Wolosker H (2006) Neuron-derived D-serine release provides a novel means to activate N-methyl-D-aspartate receptors. J Biol Chem 281:14151–14162.

Kavalali ET (2015) The mechanisms and functions of spontaneous neurotransmitter release. Nat Rev Neurosci 16:5–16.

Kim H, Ahrlund-Richter S, Wang X, Deisseroth K, Carlen M (2016) Prefrontal Parvalbumin Neurons in Control of Attention. Cell 164:208–218.

Kirihara K, Rissling AJ, Swerdlow NR, Braff DL, Light GA (2012) Hierarchical organization of gamma and theta oscillatory dynamics in schizophrenia. Biol Psychiatry 71:873–880.

Korotkova T, Fuchs EC, Ponomarenko A, von Engelhardt J, Monyer H (2010) NMDA receptor ablation on parvalbumin-positive interneurons impairs hippocampal synchrony, spatial representations, and working memory. Neuron 68:557–569.

Krystal JH, Karper LP, Seibyl JP, Freeman GK, Delaney R, Bremner JD, Heninger GR, Bowers MB, Jr., Charney DS (1994) Subanesthetic effects of the noncompetitive NMDA antagonist, ketamine, in humans. Psychotomimetic, perceptual, cognitive, and neuroendocrine responses. Arch Gen Psychiatry 51:199–214.

Kutsuwada T, Sakimura K, Manabe T, Takayama C, Katakura N, Kushiya E, Natsume R, Watanabe M, Inoue Y, Yagi T, Aizawa S, Arakawa M, Takahashi T, Nakamura Y, Mori H, Mishina M (1996) Impairment of suckling response, trigeminal neuronal pattern formation, and hippocampal LTD in NMDA receptor epsilon 2 subunit mutant mice. Neuron 16:333–344.

Lahti AC, Holcomb HH, Medoff DR, Tamminga CA (1995a) Ketamine activates psychosis and alters limbic blood flow in schizophrenia. Neuroreport 6:869–872.

Lahti AC, Koffel B, LaPorte D, Tamminga CA (1995b) Subanesthetic doses of ketamine stimulate psychosis in schizophrenia. Neuropsychopharmacology 13:9–19.

Lahti AC, Weiler MA, Tamara Michaelidis BA, Parwani A, Tamminga CA (2001) Effects of ketamine in normal and schizophrenic volunteers. Neuropsychopharmacology 25:455–467.

Lane HY, Chang YC, Liu YC, Chiu CC, Tsai GE (2005) Sarcosine or D-serine add-on treatment for acute exacerbation of schizophrenia: a randomized, double-blind, placebo-controlled study. Arch Gen Psychiatry 62:1196–1204.

Lewis DA, Volk DW, Hashimoto T (2004) Selective alterations in prefrontal cortical GABA neurotransmission in schizophrenia: a novel target for the treatment of working memory dysfunction. Psychopharmacology (Berl) 174:143–150.

Lewis DA, Hashimoto T, Volk DW (2005) Cortical inhibitory neurons and schizophrenia. Nat Rev Neurosci 6:312–324.

Lewis DA, Hashimoto T, Morris HM (2008) Cell and receptor type-specific alterations in markers of GABA neurotransmission in the prefrontal cortex of subjects with schizophrenia. Neurotox Res 14:237–248.

Lewis DA, Pierri JN, Volk DW, Melchitzky DS, Woo TU (1999) Altered GABA neurotransmission and prefrontal cortical dysfunction in schizophrenia. Biol Psychiatry 46:616–626.

Li Y, Erzurumlu RS, Chen C, Jhaveri S, Tonegawa S (1994) Whisker-related neuronal patterns fail to develop in the trigeminal brainstem nuclei of NMDAR1 knockout mice. Cell 76:427–437.

Lin H, Jacobi AA, Anderson SA, Lynch DR (2016) D-Serine and Serine Racemase Are Associated with PSD-95 and Glutamatergic Synapse Stability. Front Cell Neurosci 10:34.

Lin Y, Bloodgood BL, Hauser JL, Lapan AD, Koon AC, Kim TK, Hu LS, Malik AN, Greenberg ME (2008) Activity-dependent regulation of inhibitory synapse development by Npas4. Nature 455:1198–1204.

Ling DS, Benardo LS (1995) Recruitment of GABAA inhibition in rat neocortex is limited and not NMDA dependent. J Neurophysiol 74:2329–2335.

Lodge DJ, Behrens MM, Grace AA (2009) A loss of parvalbumin-containing interneurons is associated with diminished oscillatory activity in an animal model of schizophrenia. J Neurosci 29:2344–2354.

Lorrain DS, Baccei CS, Bristow LJ, Anderson JJ, Varney MA (2003) Effects of ketamine and N-methyl-D-aspartate on glutamate and dopamine release in the rat prefrontal cortex: modulation by a group II selective metabotropic glutamate receptor agonist LY379268. Neuroscience 117:697–706.

Lu W, Bushong EA, Shih TP, Ellisman MH, Nicoll RA (2013) The cell-autonomous role of excitatory synaptic transmission in the regulation of neuronal structure and function. Neuron 78:433–439.

Ma TM, Paul BD, Fu C, Hu S, Zhu H, Blackshaw S, Wolosker H, Snyder SH (2014) Serine racemase regulated by binding to stargazin and PSD-95: potential N-methyl-D-aspartate-alpha-amino-3-hydroxy-5-methyl-4-isoxazolepropionic acid (NMDA-AMPA) glutamate neurotransmission cross-talk. J Biol Chem 289:29631–29641.

Malhotra AK, Pinals DA, Adler CM, Elman I, Clifton A, Pickar D, Breier A (1997) Ketamine-induced exacerbation of psychotic symptoms and cognitive impairment in neuroleptic-free schizophrenics. Neuropsychopharmacology 17:141–150.

Marder CP, Buonomano DV (2004) Timing and balance of inhibition enhance the effect of long-term potentiation on cell firing. J Neurosci 24:8873–8884.

McHugh TJ, Blum KI, Tsien JZ, Tonegawa S, Wilson MA (1996) Impaired hippocampal representation of space in CA1-specific NMDAR1 knockout mice. Cell 87:1339–1349.

Mellios N, Huang HS, Baker SP, Galdzicka M, Ginns E, Akbarian S (2009) Molecular determinants of dysregulated GABAergic gene expression in the prefrontal cortex of subjects with schizophrenia. Biol Psychiatry 65:1006–1014.

Miller RF (2004) D-Serine as a glial modulator of nerve cells. Glia 47:275–283.

Miya K, Inoue R, Takata Y, Abe M, Natsume R, Sakimura K, Hongou K, Miyawaki T, Mori H (2008) Serine racemase is predominantly localized in neurons in mouse brain. J Comp Neurol 510:641–654.

Moghaddam B, Adams B, Verma A, Daly D (1997) Activation of glutamatergic neurotransmission by ketamine: a novel step in the pathway from NMDA receptor blockade to dopaminergic and cognitive disruptions associated with the prefrontal cortex. J Neurosci 17:2921–2927.

Mohn AR, Gainetdinov RR, Caron MG, Koller BH (1999) Mice with reduced NMDA receptor expression display behaviors related to schizophrenia. Cell 98:427–436.

Morita Y, Ujike H, Tanaka Y, Otani K, Kishimoto M, Morio A, Kotaka T, Okahisa Y, Matsushita M, Morikawa A, Hamase K, Zaitsu K, Kuroda S (2007) A genetic variant of the serine racemase gene is associated with schizophrenia. Biol Psychiatry 61:1200–1203.

Morris HM, Hashimoto T, Lewis DA (2008) Alterations in somatostatin mRNA expression in the dorsolateral prefrontal cortex of subjects with schizophrenia or schizoaffective disorder. Cereb Cortex 18:1575–1587.

Mothet JP, Parent AT, Wolosker H, Brady RO, Jr., Linden DJ, Ferris CD, Rogawski MA, Snyder SH (2000) D-serine is an endogenous ligand for the glycine site of the N-methyl-D-aspartate receptor. Proc Natl Acad Sci U S A 97:4926–4931.

Moy SS, Perez A, Koller BH, Duncan GE (2006) Amphetamine-induced disruption of prepulse inhibition in mice with reduced NMDA receptor function. Brain Res 1089:186–194.

Moy SS, Nikolova VD, Riddick NV, Baker LK, Koller BH (2012) Preweaning sensorimotor deficits and adolescent hypersociability in Grin1 knockdown mice. Dev Neurosci 34:159–173.

Nakazawa K, Sapkota K (2020) The origin of NMDA receptor hypofunction in schizophrenia. Pharmacol Ther 205:107426.

Nakazawa K, Jeevakumar V, Nakao K (2017) Spatial and temporal boundaries of NMDA receptor hypofunction leading to schizophrenia. NPJ Schizophr 3:7.

Panatier A, Theodosis DT, Mothet JP, Touquet B, Pollegioni L, Poulain DA, Oliet SH (2006) Glia-derived D-serine controls NMDA receptor activity and synaptic memory. Cell 125:775–784.

Quintero GC, Erzurumlu RS, Vaccarino AL (2008) Evaluation of morphine analgesia and motor coordination in mice following cortex-specific knockout of the N-methyl-D-aspartate NR1-subunit. Neurosci Lett 437:55–58.

Ramsey AJ (2009) NR1 knockdown mice as a representative model of the glutamate hypothesis of schizophrenia. Prog Brain Res 179:51–58.

Rosoklija G, Toomayan G, Ellis SP, Keilp J, Mann JJ, Latov N, Hays AP, Dwork AJ (2000) Structural abnormalities of subicular dendrites in subjects with schizophrenia and mood disorders: preliminary findings. Arch Gen Psychiatry 57:349–356.

Saunders JA, Gandal MJ, Siegel SJ (2012) NMDA antagonists recreate signal-to-noise ratio and timing perturbations present in schizophrenia. Neurobiol Dis 46:93–100.

Saunders JA, Tatard-Leitman VM, Suh J, Billingslea EN, Roberts TP, Siegel SJ (2013) Knockout of NMDA receptors in parvalbumin interneurons recreates autism-like phenotypes. Autism Res 6:69–77.

Schell MJ, Molliver ME, Snyder SH (1995) D-serine, an endogenous synaptic modulator: localization to astrocytes and glutamate-stimulated release. Proc Natl Acad Sci U S A 92:3948–3952.

Schell MJ, Brady RO, Jr., Molliver ME, Snyder SH (1997) D-serine as a neuromodulator: regional and developmental localizations in rat brain glia resemble NMDA receptors. J Neurosci 17:1604–1615.

Shi J, Badner JA, Gershon ES, Liu C (2008) Allelic association of G72/G30 with schizophrenia and bipolar disorder: a comprehensive meta-analysis. Schizophr Res 98:89–97.

Singer W, Gray CM (1995) Visual feature integration and the temporal correlation hypothesis. Annu Rev Neurosci 18:555–586.

Sohal VS (2016) How Close Are We to Understanding What (if Anything) gamma Oscillations Do in Cortical Circuits? J Neurosci 36:10489–10495.

Sohal VS, Rubenstein JLR (2019) Excitation-inhibition balance as a framework for investigating mechanisms in neuropsychiatric disorders. Mol Psychiatry 24:1248–1257.

Stan AD, Lewis DA (2012) Altered cortical GABA neurotransmission in schizophrenia: insights into novel therapeutic strategies. Curr Pharm Biotechnol 13:1557–1562.

Steullet P, Cabungcal JH, Coyle J, Didriksen M, Gill K, Grace AA, Hensch TK, LaMantia AS, Lindemann L, Maynard TM, Meyer U, Morishita H, O’Donnell P, Puhl M, Cuenod M, Do KQ (2017) Oxidative stress-driven parvalbumin interneuron impairment as a common mechanism in models of schizophrenia. Mol Psychiatry 22:936–943.

Suzuki Y, Jodo E, Takeuchi S, Niwa S, Kayama Y (2002) Acute administration of phencyclidine induces tonic activation of medial prefrontal cortex neurons in freely moving rats. Neuroscience 114:769–779.

Szabadits E, Cserep C, Szonyi A, Fukazawa Y, Shigemoto R, Watanabe M, Itohara S, Freund TF, Nyiri G (2011) NMDA receptors in hippocampal GABAergic synapses and their role in nitric oxide signaling. J Neurosci 31:5893–5904.

Taniguchi H, He M, Wu P, Kim S, Paik R, Sugino K, Kvitsiani D, Fu Y, Lu J, Lin Y, Miyoshi G, Shima Y, Fishell G, Nelson SB, Huang ZJ (2011) A resource of Cre driver lines for genetic targeting of GABAergic neurons in cerebral cortex. Neuron 71:995–1013.

Tatard-Leitman VM, Jutzeler CR, Suh J, Saunders JA, Billingslea EN, Morita S, White R, Featherstone RE, Ray R, Ortinski PI, Banerjee A, Gandal MJ, Lin R, Alexandrescu A, Liang Y, Gur RE, Borgmann-Winter KE, Carlson GC, Hahn CG, Siegel SJ (2015) Pyramidal cell selective ablation of N-methyl-D-aspartate receptor 1 causes increase in cellular and network excitability. Biol Psychiatry 77:556–568.

Tiesinga P, Sejnowski TJ (2009) Cortical enlightenment: are attentional gamma oscillations driven by ING or PING? Neuron 63:727–732.

Tiesinga PH, Fellous JM, Salinas E, Jose JV, Sejnowski TJ (2004) Inhibitory synchrony as a mechanism for attentional gain modulation. J Physiol Paris 98:296–314.

Ting JT, Lee BR, Chong P, Soler-Llavina G, Cobbs C, Koch C, Zeng H, Lein E (2018) Preparation of Acute Brain Slices Using an Optimized N-Methyl-D-glucamine Protective Recovery Method. J Vis Exp.

Tsai G, Yang P, Chung LC, Lange N, Coyle JT (1998) D-serine added to antipsychotics for the treatment of schizophrenia. Biol Psychiatry 44:1081–1089.

Tsien JZ, Huerta PT, Tonegawa S (1996) The essential role of hippocampal CA1 NMDA receptor-dependent synaptic plasticity in spatial memory. Cell 87:1327–1338.

Ultanir SK, Kim JE, Hall BJ, Deerinck T, Ellisman M, Ghosh A (2007) Regulation of spine morphology and spine density by NMDA receptor signaling in vivo. Proc Natl Acad Sci U S A 104:19553–19558.

van den Buuse M (2010) Modeling the positive symptoms of schizophrenia in genetically modified mice: pharmacology and methodology aspects. Schizophr Bull 36:246–270.

Vollenweider FX, Leenders KL, Scharfetter C, Antonini A, Maguire P, Missimer J, Angst J (1997) Metabolic hyperfrontality and psychopathology in the ketamine model of psychosis using positron emission tomography (PET) and [18F]fluorodeoxyglucose (FDG). Eur Neuropsychopharmacol 7:9–24.

Wigstrom H, Gustafsson B (1983) Facilitated induction of hippocampal long-lasting potentiation during blockade of inhibition. Nature 301:603–604.

Williams JH, Kauer JA (1997) Properties of carbachol-induced oscillatory activity in rat hippocampus. J Neurophysiol 78:2631–2640.

Wolosker H, Blackshaw S, Snyder SH (1999) Serine racemase: a glial enzyme synthesizing D-serine to regulate glutamate-N-methyl-D-aspartate neurotransmission. Proc Natl Acad Sci U S A 96:13409–13414.

Wolosker H, Panizzutti R, De Miranda J (2002) Neurobiology through the looking-glass: D-serine as a new glial-derived transmitter. Neurochem Int 41:327–332.

Wolosker H, Balu DT, Coyle JT (2016) The Rise and Fall of the d-Serine-Mediated Gliotransmission Hypothesis. Trends Neurosci 39:712–721.

Wong JM, Gray JA (2018) Long-Term Depression Is Independent of GluN2 Subunit Composition. J Neurosci 38:4462–4470.

Wong JM, Folorunso O, Barragan EV, Berciu C, Harvey TL, DeChellis-Marks MR, Glausier JR, MacDonald ML, Coyle JT, Balu DT, Gray JA (2020) Postsynaptic serine racemase regulates NMDA receptor function. bioRxiv:2020.2006.2016.155572.

Yoshikawa M, Takayasu N, Hashimoto A, Sato Y, Tamaki R, Tsukamoto H, Kobayashi H, Noda S (2007) The serine racemase mRNA is predominantly expressed in rat brain neurons. Arch Histol Cytol 70:127–134.

Zerucha T, Stuhmer T, Hatch G, Park BK, Long Q, Yu G, Gambarotta A, Schultz JR, Rubenstein JL, Ekker M (2000) A highly conserved enhancer in the Dlx5/Dlx6 intergenic region is the site of cross-regulatory interactions between Dlx genes in the embryonic forebrain. J Neurosci 20:709–721.

